# PINK1-G411S mutant increases kinase stability and enhances mitochondrial-linked functions

**DOI:** 10.1101/2024.06.28.601304

**Authors:** Filipa B. Gonçalves, Francisco J. Enguita, Vanessa A. Morais

## Abstract

PINK1, a mitochondria targeted Serine/Threonine kinase, regulates ATP production by phosphorylating the Complex I subunit NdufA10. However, when in the presence of depolarized mitochondria, PINK1 phosphorylates ubiquitin and Parkin triggering mitochondria clearance. Mutations in PINK1 have been linked to early-onset recessive familial forms of Parkinson’s disease (PD). Deficits in Complex I enzymatic activity and an increase in oxidative damage have been identified in multiple brain regions of PD patients. Unravelling how PINK1 activity regulates mitochondria fate is pivotal. In the present study we characterized how human PD-related PINK1 mutants affect major PINK1 functions. Using molecular dynamics, we gain mechanistic insight into how specific mutations alter the tertiary structure and stability of PINK1’s ATP-binding pocket, leading to an increased rigidity and stability. More importantly, we report a structural explanation for the enhanced kinase function of the PINK1-G411S mutant.

## INTRODUCTION

Parkinson’s disease (PD), the second most common neurodegenerative disorder, is characterized by loss of dopaminergic neurons in the substantia nigra (SN) pars compacta and the presence of Lewy bodies(1). A link between PD and mitochondrial dysfunctions was established when the mitochondrial toxin metabolite of 1-Methyl-4-phenyl-1,2,3,6-tetrahydropyridine (MPTP), MPP+, was found to inhibit the electron transport chain Complex I(2) in humans. Later, mutations in the mitochondria associated genes PTEN-induced putative kinase-1 (PINK1)(3) and Parkin(4), amongst other genes(5), were found in familial autosomal recessive forms of PD. These findings further strengthened the link between mitochondrial homeostasis and PD.

PINK1, a nuclear encoded gene, is a 581 amino acid long serine/threonine kinase with a N-terminal mitochondria targeting sequence(3) and has been linked to many different cellular pathways(6). Under basal conditions, PINK1 is imported into the mitochondria, where it is firstly processed by the mitochondrial processing peptidase (MPP), a matrix protease which cleaves the N-terminal peptides from precursor proteins, and subsequently by the Presenilin-associated rhomboid-like protein (PARL)(7–10). Some reports have shown that this PARL-mediated PINK1 cleavage product is translocated to the cytoplasm where it is degraded in a proteasome dependent manner(11). This rapid turnover results in low levels of endogenous PINK1(12). When mitochondria are depolarized, PINK1 mitochondrial import is impaired and PINK1 is stabilized on the mitochondrial outer membrane where in combination with Parkin triggers the clearance of damaged mitochondria via the mitophagy pathway(13–15). PINK1 initiates mitophagy by phosphorylating Ubiquitin(16) and Parkin(17–19). For this, PINK1 needs to dimerize and be autophosphorylated on residues Ser228 and Ser402(20,21).

To date, the Human Gene Mutation Database (HGMB) has reported at least 130 homozygous or compound heterozygous mutations in the PINK1 gene. Most of these mutations result in loss of function of this gene and are a well-known cause of early-onset PD(22). PINK1 patients mirror sporadic PD symptomatology, with bradykinesia, rigidity, tremors and present a good response to levodopa, and although PINK1 mutations rarely lead to cognitive decline, a higher rate of psychiatric manifestations occurs(23). Albeit quite studied, how PINK1 mutations drive loss of dopaminergic neurons and consequently PD-linked phenotypes remains unclear.

To scrutinize the impact that PINK1 mutations have on PINK1 function and overall mitochondrial homeostasis, we systematically analyzed several human PD-causing hPINK1 mutations (C92F, A168P, H271Q, G309D, L347P, G411S, E417G and L489P) (Figure 1A) and determined the impact that each mutation has on described molecular pathways and protein structure. Our results showed that most studied human mutations have an impact on PINK1 kinase activity, disrupting the mitochondrial quality control. Yet, we observed a peculiar result for PINK1-G411S, where a gain of kinase activity for the substrate Parkin is observed. Several controversial reports about the G411S mutation are present in the literature. It is either reported as preventive measure for mitochondrial quality control, by reducing PINK1 kinase activity in heterozygous carriers(24) or not even considered as a major risk factor or causative for PD in heterozygous variants(25). Regardless, PINK1-G411S variation is more prevalent in Norwegian, Polish and Swedish population(24,26,27), and there is no clear common pattern of clinical characteristics. Following our data, and due to this knowledge gap and controversial theories, we decided to further investigate the PINK1-G411S mutation. Here, we report that the human mutant PINK1-G411S harbours an additional phosphorylation site. Also, molecular modelling and dynamic simulations suggest that the additional phosphorylation site can induce a conformational stabilization of the ATP-binding pocket of the enzyme, characterized by a pattern of tighter molecular interactions across the functional amino acid residues. These findings can explain the enhanced enzymatic function of the G411S mutant towards the function of PINK1 in mitochondrial quality control.

**Figure 1.**
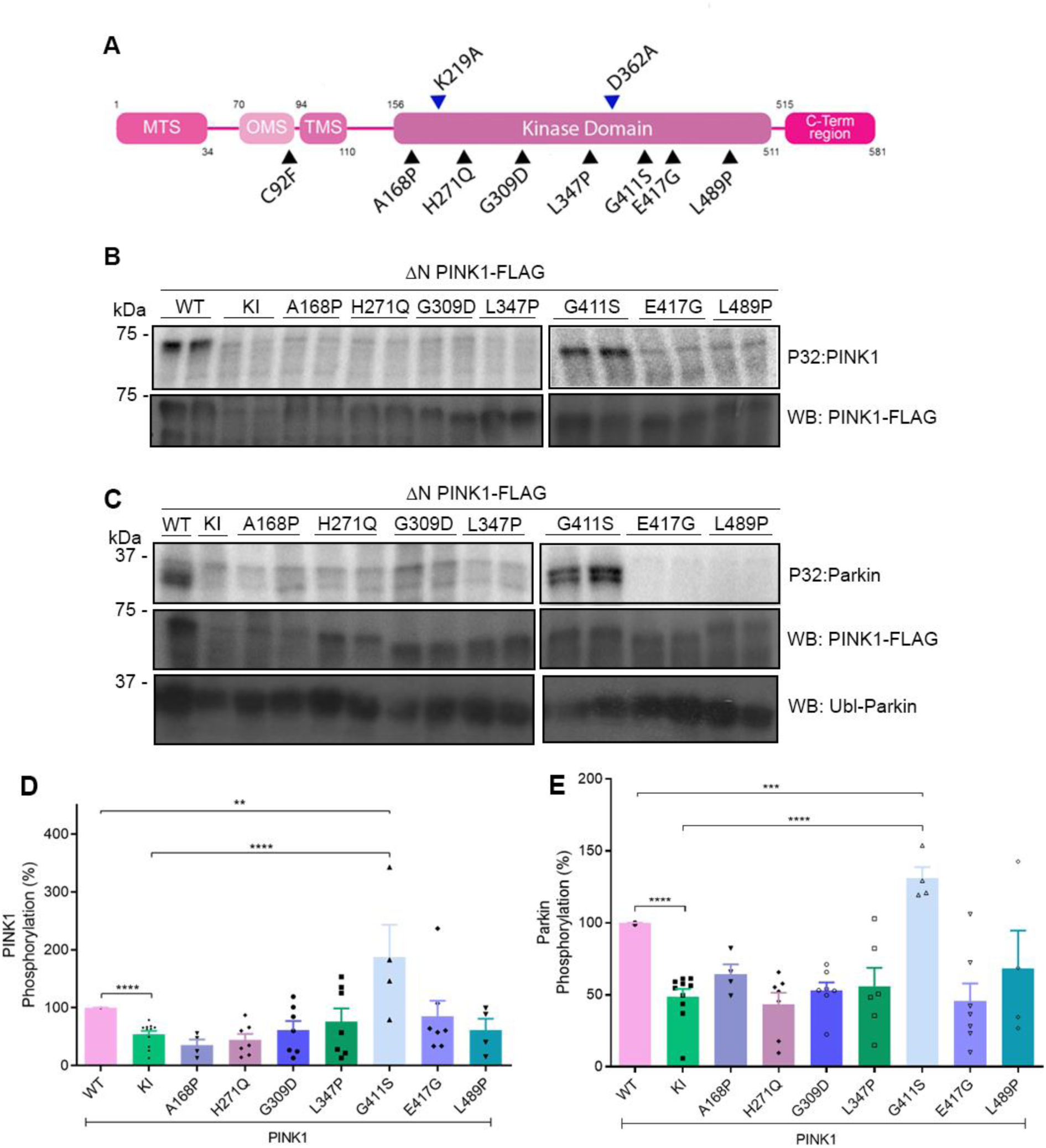
Impact of hPINK1 mutants in unhealthy mitochondria pathways. Human PINK1 domains and studied mutations. PINK1 is divided in different regions, at N-terminal region: mitochondrial targeting sequence (MTS), mitochondria membrane localization (OMS) and transmembrane sequence (TMS). The majority of the studied mutations reside in the kinase domain, with the only exception of C92F, which is in the OMS. In this study a kinase inactive control was used, which contains the two mutations signed in blue (K219A and D362A, responsible for ATP positioning and catalytic loop, respectively). B), C) An in vitro phosphorylation assay using [γ-32P]-ATP was performed with purified WT and KI PINK1-FLAG and purified GST-Ubl Parkin as substrate, respectively. Autoradiogram phosphorlation levels were normalized to total amount of PINK1-FLAG prontein. D), E) Quantification of [γ-32P]-ATP in vitro PINK1 autophosphorylation (D) and phosphorylation fo GST-Parkin (E). All studied mutations are significantly different from PINK1-WT, presenting phosphorylation levels more comparable to PINK1-KI; with the exception of PINK1-G411S that is statistical increased comparing to both controls, PINK1-KI and -WT. Statistical significance was calculated using Student’s t-test (mean +/- SEM, minimal n=4 independent experiments; * P < 0.05; ** P < 0.01; *** P < 0.001; **** P < 0.0001; n.s. non-significant).

## RESULTS

### PINK1-G411S mutant phosphorylates PINK1 and Parkin

The PINK1 gene encodes for a 581 amino acids long nuclear encoded protein, known to be mutated in familiar forms of PD. In this study we decided to focus on 8 described human missense mutations (Figure 1A), based on their pathogenicity, role as a risk factor in PD and a transmembrane residing mutation. Although PINK1 insect orthologues have been widely used to study PINK1 function, several crucial differences in the catalytic activity(28,29) and overall structure(30,31) have been reported. Taking these differences into account, and aiming to understand the biological relevance of PD-linked PINK1 mutations, we pursued this study using the human form of PINK1. The majority of human PINK1 mutations are located within the kinase domain, demonstrating the importance of PINK1 kinase activity in PD pathogenesis(6). For this study, the mutations A168P, H271Q, G309D, L347P, G411S, E417G and L489P were selected to represent the mutations within the kinase domain. Moreover, G309D has been widely reported as a loss of kinase activity mutation(32). The Leu to Pro mutants, L347P and L489P, have already been correlated with disturbed protein stability(30–33). The A168P mutation is reported to influence the overall kinase structure(34). Both G411S and E417G are located in PINK1 activation loop, which is crucial for dimer structure formation(30,31). The C92F mutation was selected to determine the function of the putative hydrophobic patch (transmembrane sequence, TMS) and the N-terminal (Figure 1A).

When PINK1/Parkin mediated mitophagy is triggered, the PINK1-mediated phosphorylation of PINK1 itself and Parkin is one of the initial and fundamental steps of this pathway(14,15). In order to assess whether PINK1 PD-related mutants affect the ability of this kinase to phosphorylate PINK1 and its substrate Parkin, an in vitro phosphorylation assay was implemented(21). Incorporation of radiolabeled phosphate was assessed via autoradiogram and results were normalized to PINK1 expression levels (Figure 1B, C). As previous described(21), we also detected an absence of in vitro autophosphorylation of full-length (FL) PINK1, therefore a truncated form of PINK1 lacking amino acids 1-113 was used. As expected, PINK1-WT is able to phosphorylate itself (Figure 1B) and Parkin (Figure 1C).

On the other hand, PINK1 kinase inactive (KI) form was unable to phosphorylate either substrate (Figure 1D, E). In agreement with previous reports, the mutants A168P, H271Q, G309D, L347P and E417G were not able to induce phosphorylation(20,32). Interestingly, the PINK1-G411S form was the only mutant able to phosphorylate both PINK1 and Parkin to significantly higher levels than those obtained for PINK1-WT (Figure 1D, E), suggesting that this mutation may increase the overall kinase activity of PINK1.

### PINK1-G411S mutant is able to induce mitophagy

Although the studied PINK1 mutant forms are unable to phosphorylate PINK1 itself and Parkin, and as Parkin recruitment to damaged mitochondria may precede these phosphorylation events(35–37), we determined their ability towards Parkin recruitment and sequential induction of mitophagy. For this, HeLa WT and HeLa PINK1^-/-^ (Figure 2A, Figure S1C-H) cells were transfected with PINK1 mutant forms and with GFP-tagged-Parkin, as previously described(21) (Figure S1C-H). In basal conditions (DMSO), Parkin is predominately located in the cytosol and does not co-localize with mitochondria (Figure 2A). However, when PINK1 is re-introduced in HeLa PINK1^-/-^ cells depolarized with CCCP, a mitochondrial uncoupler, a fragmented mitochondrial pattern is observed, and Parkin is recruited to the mitochondria co-localizing with the mitochondrial marker HSP60 (Figure 2A, Figure S1D). This is not observed in HeLa PINK1^-/-^ cells treated with CCCP, in the absence of PINK1 or in the presence of a PINK1-KI, thus Parkin is not recruited to mitochondria (Figure 2C, Figure S1D). However, in HeLa WT background when PINK1-KI is expressed a reduced recruitment of Parkin to mitochondria is observed (Figure 2B), as reported in *Drosophila*(36), other HeLa cells studies(37), PINK1 KO cortical neurons(37) and mouse embryonic fibroblast (MEF) cell lines(38), indicating that PINK1 has a crucial role in Parkin recruitment to mitochondria.

**Figure 2.**
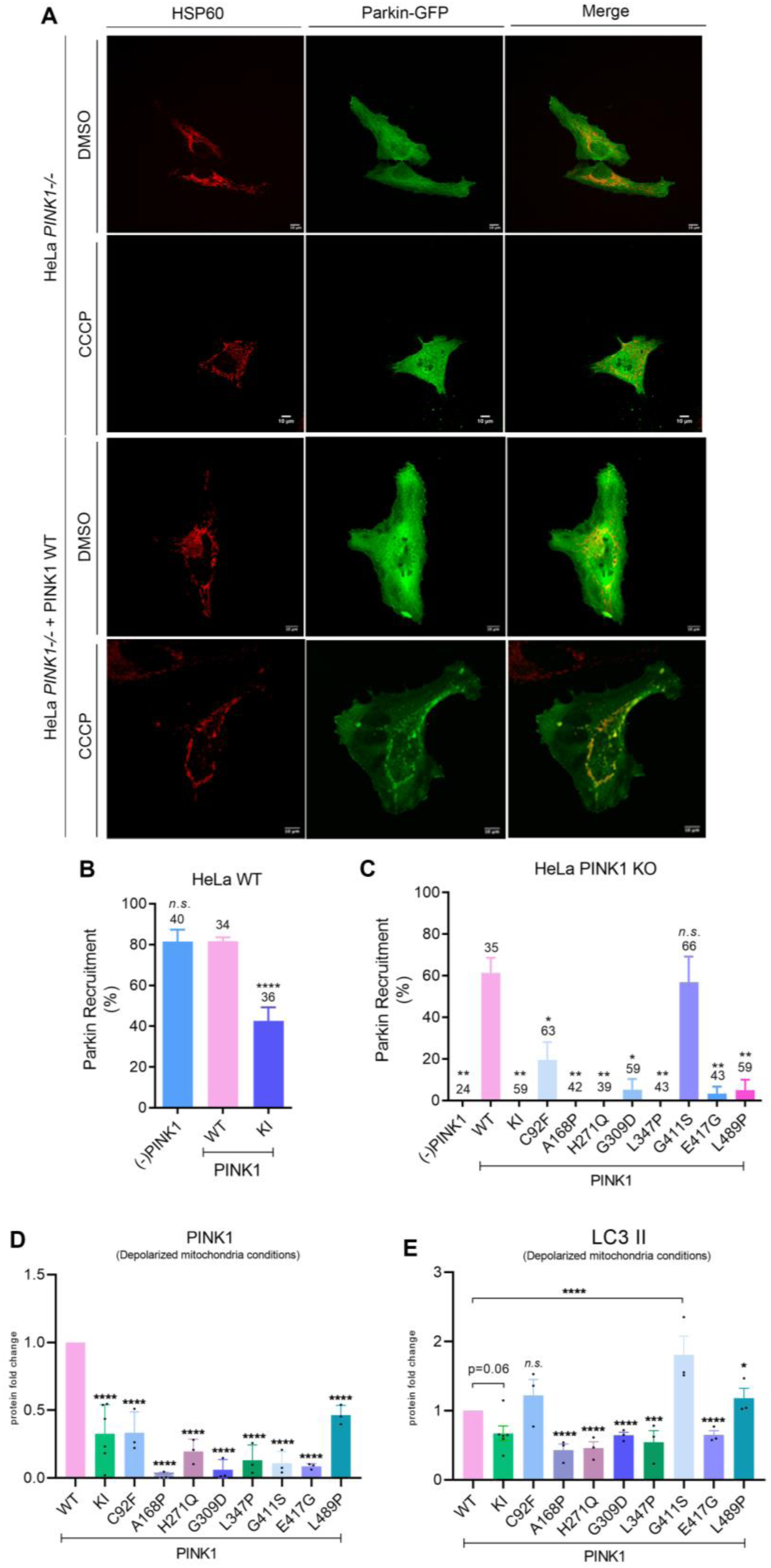
Impact of PINK1 mutations in mitochondrial quality control. A) HeLa PINK1^-/-^ cells were co-transfected with Parkin-GFP and PINK1-WT and treated with DMSO or 10µM CCCP for 3hours, to induce depolarization. In DMSO conditions there is a network of mitochondria (HSP60) spread throughout the cell, which is disrupted when treated with CCCP. Also, in CCCP conditions Parkin presents a pattern similar to the mitochondrial marker HSP60, indicating that Parkin was recuited to mitochondria in this condition (Scale bar=10µm). This Parkin recruitment pattern is not observed in the PINK1^-/-^ conditions. B), C) Quantification of Parkin recruited to mitochondria in HeLa PINK1^-/-^ cells (-PINK1) and HeLa WT cells expressing PINK1-WT or PINK1-KI, and all cells were co-transfected with Parkin-GFP (B); and HeLa PINK1^-/-^ cells co-transfected with Parkin-GFP and PINK1-WT, -KI or cliical mutants (C). Parkin recruitment only occurs at levels comparable to what is observed in HeLa WT when PINK1-G411S mutant is present. Pecentage of Parkin recruitment is compared, in each cell line, to PINK1-WT transfection. Cell counts are represented above each corresponding bar. D), E) Western Blot quantification for the expression levels of PINK1 forms (D) and LC3 II (E) in MEF cells treated with DMSO or 25µM of CCCP. Both, PINK1 and LC3 II levels were normalized to GAPDH. Statistical significance was calculated using Student’s t-test (mean +/- SEM, n=3 independent experiments; * P < 0.05; ** P < 0.01; *** P < 0.001; **** P < 0.0001; n.s. non-significant).

As also observed in the phosphorylation assay, in this Parkin recruitment assay, the PINK1-G411S is the only mutant able to restore Parkin recruitment to mitochondria to levels comparable with PINK1-WT, which corroborates what was already shown in MEF cells(38) (Figure 2C). Also, the PINK1-C92F can recruit Parkin to mitochondria at a lower extent (Figure 2C, Figure S1E).

To assess the induction of the mitophagy/autophagy pathway, especially in the presence of the PINK1-G411S mutant, we analyzed the accumulation of PINK1 to mitochondria and the detection of LC3-II, which is required to initiate formation and lengthening of the autophagosome(39). For this, MEF cells stably transduced with PINK1 mutants were depolarized with CCCP and protein levels were assessed (Figure 2D, E; Figure S2A). In depolarization conditions (CCCP+), PINK1 levels were lower in cells expressing the PINK1 mutant forms, however LC3-II levels were significantly increased in the PINK1-G411S mutant when compared to PINK1-WT (Figure 2D-E). LC3-II increase was confirmed via immunofluorescence (Figure S2B-C), where decreased colocalization of LC3 with mitochondria in PINK1-/- was confirmed (Figure S2D). Colocalization was restored when PINK1-WT or mutant forms were re-introduced (Figure S2D). However, a clear difference between genotypes was observed when LC3 puncta area/cell was calculated (Figure S2E), where in MEF PINK1-/- or in the case of the re-introduction of PINK1-KI or PINK1-G309D mutant a clear reduction in LC3 puncta number occurred (Figure S2E), while once again PINK1-G411S has a significant increase.

Even though we analyzed several PINK1 mutant forms, intriguingly, the PINK1-G411S mutant can restore the PINK1 phenotype by phosphorylating truncated PINK1 and Parkin, and by triggering mitochondria for mitophagy in the presence of depolarized mitochondria.

### Absence of PINK1 induces mitochondrial fragmentation and mass loss

Although still a matter of debate(40,41)] absence of PINK1 has been associated with alterations in overall mitochondrial morphology(18,42–46). To assess mitochondria morphology, we transiently expressed a mitochondrial targeted DsRed tagged expression vector (Mito-DsRed) in MEF cells stably transduced with PINK1-WT, PINK1-KI or PINK1 mutant forms. We observed a clear increase of fragmented mitochondria in a PINK1^-/-^ phenotype, whereas endogenously expressed PINK1 presents higher levels of elongated mitochondria (Figure 3A, B).

**Figure 3.**
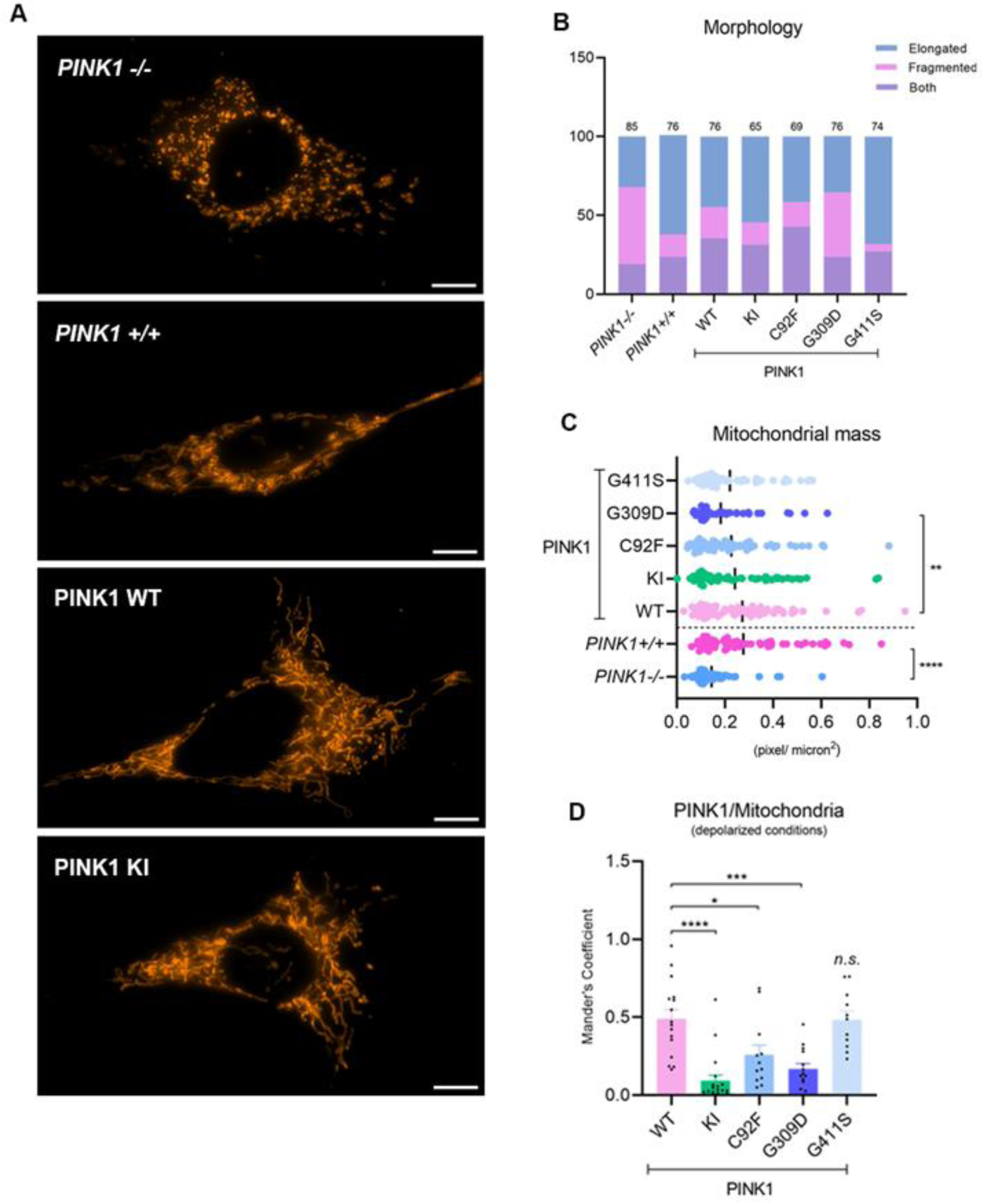
Impact of PINK1 mutants in mitochondria morphology. A) MEF cell line endogenously expressing PINK1, or transduced with PINK1-WT, -KI and mutant forms, transiently expressing plasmid MitodsRed (Scale bar=10µm) and B) consequent mitochondrial morphology analysis, showing a clear increased on fragmented mitochondrial in PINK1^-/-^, andC) Mitochondrial mass was evaluated in MEF cell line endogenously expressing PINK1 mutant forms using ImageJ software. The cell area was calculated and mitochondria corresponding pixels were accessed (only significant differences were highlighted) D) PINK1 colocalization with Mitochondria was accessed in HeLa PINK1^-/-^ cells, transfected with PINK1-FLAG tagged constructs. Cells were treated for 3hours with 10µM of CCCP. Colocalization was accessed via ImageJ plug-in JACoP which calculates Mander’s Coefficient, which diverges between 0 to 1, the prior corresponding to non-overlapping images and the last to 100% colocalization. Statistical significance was calculated using one way ANOVA followed by Dunnets test (mean +/- SEM, n=3 independent experiments; * P < 0.05; ** P < 0.01; *** P < 0.001; **** P < 0.0001; n.s. – non significant).

Morphology does not seem to be affected by the lack of kinase activity of PINK1 (Figure 3A, B). Interestingly, the PINK1-G309D mutant has a higher presence of fragmented mitochondria, while PINK1-G411S mutant has the opposite phenotype (Figure 3A, B; Figure S3A). Interestingly, mitochondrial mass follows the same pattern, where PINK1^-/-^ and PINK1-G309D have a significant decrease of mitochondrial mass (Figure 3C). We aimed to understand if observed alterations in mitochondria morphology and mass could impact PINK1 presence in mitochondria upon depolarization, for this we transfected HeLa PINK1^-/-^ cells, with FLAG tagged PINK1-WT and –KI, as well as, the PINK1 mutants C92F, G309D and G411S and treated cells with 10µM CCCP for 3hours. We observed that the kinase inactive form of PINK1 and the mutant PINK1-G309D have a significantly reduced localization with mitochondria (Figure 3D; Figure S3B, C). Considering that PINK1 dimerization could be dependent on PINK1’s phosphorylation capability(31,35), one could assume that PINK1-KI has impaired mitochondria-PINK1 colocalization, as well as the mutations with an impaired PINK1 kinase activity (Figure 1D, E). Overall, and only considering the PINK1 mutants analyzed in this study, PINK1 contributes to mitochondrial integrity, and this is only disrupted when PINK1-G309D mutant is present.

### Presence and activity of PINK1 is needed for Complex I activity

The complexes of the Electron Transport Chain (ETC) are responsible for maintaining the mitochondrial membrane potential, by pumping protons, resultant from the oxidation of NADH and FADH_2_, from the matrix into the intermembrane space. The generated proton gradient drives the ATP production at complex V (F1F0-ATP synthase). Nonetheless, if oxidative phosphorylation is impaired, Complex V can pump protons in the reverse direction, at the expense of hydrolyzing ATP molecules, hence, maintaining the mitochondrial membrane potential. PINK1 deficiency has an impact on the enzymatic function of Complex I, causing a decrease in membrane potential(40). We analyzed the enzymatic activity of individual OXPHOS complexes in mitochondrial enriched fractions from MEF cells harboring PINK1 mutations (Figure 4A, B). We confirmed that both presence and activity of PINK1-WT is required for a functional Complex I (NADH:ubiquinone oxidoreductase) (Figure 4A). Curiously, Complex I activity seems to be unaffected in PINK1-C92F and PINK1-G411S mutants, whereas PINK1-KI and PINK1-G309D leads to a decrease in Complex I activity. Interestingly, when looking into Complex V activity (ATPase reductase), the absence of PINK1 or the presence of PINK1-KI did not impact the activity of Complex V (Figure 4B). However, the presence of PINK1-C92F, - G309D, and -G411S significantly decreased the enzymatic activity of Complex V, suggesting an impairment in ETC. The protein expression for both Complex I and Complex V was accessed (Figure 4C-E; Figure S4). For Complex V, no significant differences in protein levels were observed (Figure 4E), indicating that differences observed in activity are not due to reduced protein levels but to overall enzymatic function. For Complex I, differences in protein levels were observed (Figure 4C, D; Figure S4). We postulate that these differences in Complex I subunits expression and functional impairment could be due to complex misassembling, consequent of oxidatively damaged Complex I core and accessory subunits, as reported in PD patient brain samples(47).

**Figure 4.**
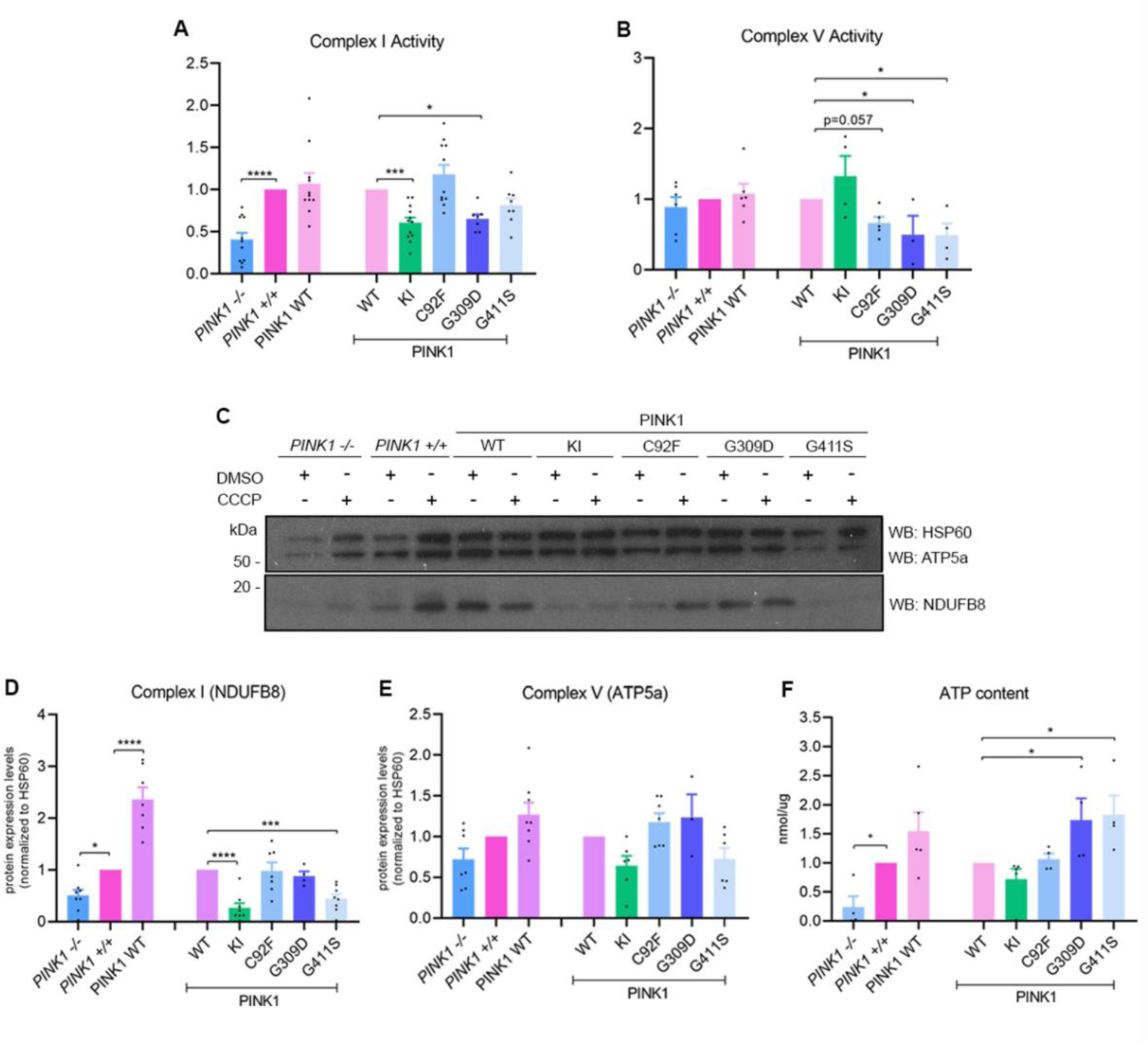
The effect of PD-related PINK1 mutations on OXPHOS. A), B) Respiratory chain measurements were performed on mitochondrial enriched fractions from MEF PINK1^+/+^ and PINK1^-/-^, and also from stably transduced MEF cells expressing PINK1-WT, PINK1-KI and respective PINK1 mutant forms. Spectrophotometric assays for the activity of Complex I (NADH:ubiquinone oxidoreductase, rotenone sensitive) (A) and Complex V (ATPsynthase, oligomycin sensitive) (B) and citrate synthase enzyme activities were performed. Values were plotted according to the ration between the specific complex’s activity and citrate synthase activity. C) – E) Western blot analyses and corresponding quantification of MEF cells treated with DMSO or 25µM of CCCP were performed to determine the expression levels of Complex I NDUFB8 (C, D) and Complex V ATP5a subunits (C, E) (NDUFB8 and ATP5a are part of the OXPHOS antibody cocktail). F) ATP content was measured in lysates from MEF PINK1^+/+^ and PINK1^-/-^ cells, and also from stably transduced MEF cells expressing PINK1-WT, -KI and respective PINK1 mutant forms. Statistical significance was calculated using one way ANOVA followed by Dunnets test (mean +/- SD, n=3 independent experiments, values for each replicate within the independent experiment is depicted in the graph; * P < 0.05; ** P < 0.01; *** P < 0.001; **** P < 0.0001; n.s. – non significant).

As lack of PINK1 impacts Complex I and PINK1 mutations impair Complex V enzymatic activity, we decided to assess ATP content. As previously described(17,18), the levels of cellular ATP are reduced in the absence of PINK1 (Figure 4F). However, in the presence of PINK1-G309D and PINK1-G411S mutants a significant increase in ATP content was observed. This can be explained by the decrease in ATPase reductase activity (Figure 4B) and may indicate differences in the overall bioenergetics profile of cells harboring these mutations.

### PINK1 absence instigates metabolic changes

To investigate if absence of PINK1 or PINK1 mutants could also disrupt further mitochondria related metabolic parameters, we explored mitochondrial respiration as a quantitative metric of mitochondrial function via OXPHOS. MEF PINK1^+/+^ and PINK1^-/-^ cells or cells with re-introduced human PINK1 mutant forms were exposed to respiration modulators and real-time measurement of changes in oxygen (oxygen consumption rate - OCR) and proton concentrations (extracellular acidification rate - ECAR) in the extracellular medium were accessed (Figure 5A; Figure S5A). Interestingly, we could observe that PINK1^-/-^ presents a higher maximum respiration when compared to PINK1^+/+^ (Figure 5C), indicating a higher flexibility to respond to an energy demands. Which seems to be a compensatory effect to the absence of PINK1, since neither PINK1-KI nor mutant forms disrupt any of the studied parameters (Figure 5B-F). ECAR, which is the acidification of the extracellular medium and can be correlated with lactate production (glycolysis), does not seem to be affected (Figure S5A), but when we look at the energy map, PINK1^+/+^ are more quiescent, whereas PINK1^-/-^ are more energetic, relying more on OXPHOS (Figure S5B-C), especially upon stress conditions. On the other hand, PINK1-KI does not have a significant metabolic potential, since they do not present a high shift when stressed. All mutations (Figure S5D) follow this pattern, however, present an even lower capability to deal with stress response than any of the controls (Figure S5C, D).

**Figure 5.**
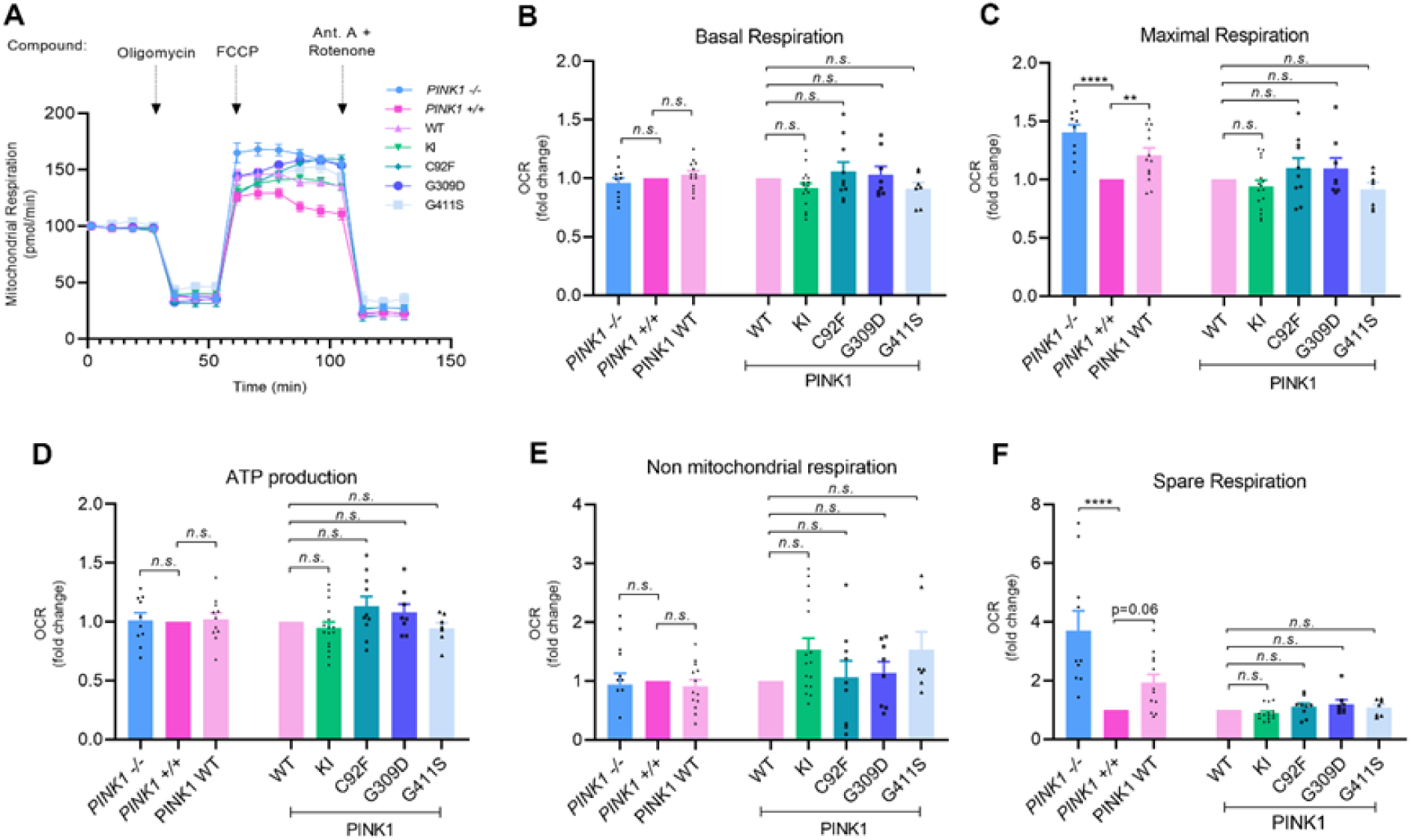
The effect of PD-related PINK1 mutations on mitochondrial bioenergetics. A) Respiration rates of MEF PINK1^+/+^, PINK1^-/-^ and stably transducted PINK1 mutants in response to bioenergetic modulators was determined. MEF cells were treated with Oligomycin, FCCP and combination of Rotenone + Antimycin A. Oligomycin inhibits respiration by inhibiting ATP synthase, while FCCP stimulates mitochondrial respiration by uncoupling ATP synthesis, while combination of Rotenone and Antimycin A shuts down mitochondrial respiration. B) - F) Through OCR responses to modulators several mitochondrial parameters were calculated: Basal Respiration (B), Maximal Respiration (C), ATP production (D), Non-mitochondrial Respiration (E) and Spare Respiration (E). Statistical significance was calculated using one way ANOVA followed by Dunnets test (mean +/- SEM, n=3 independent experiments; * P < 0.05; ** P < 0.01; *** P < 0.001; **** P < 0.0001; n.s. – non significant).

### PINK1-G411S presents a new phosphorylation site

PINK1-G411S mutation, results from an aminoacid change from a Glycine to a Serine at position 411. Glycine is a very stable aminoacid with a single hydrogen atom in its chain. Due to its compact form, it is frequently located within alpha-helices in secondary protein structure. Serine is a polar aminoacid harboring a hydroxymethyl group as a side chain. Considering characteristics of aminoacid change present at PINK1-G411S mutation site (Figure 6A) and results obtained in this work (Figure 1 & Figure 2), we postulate that this mutation could lead to an additional phosphorylation site.

**Figure 6.**
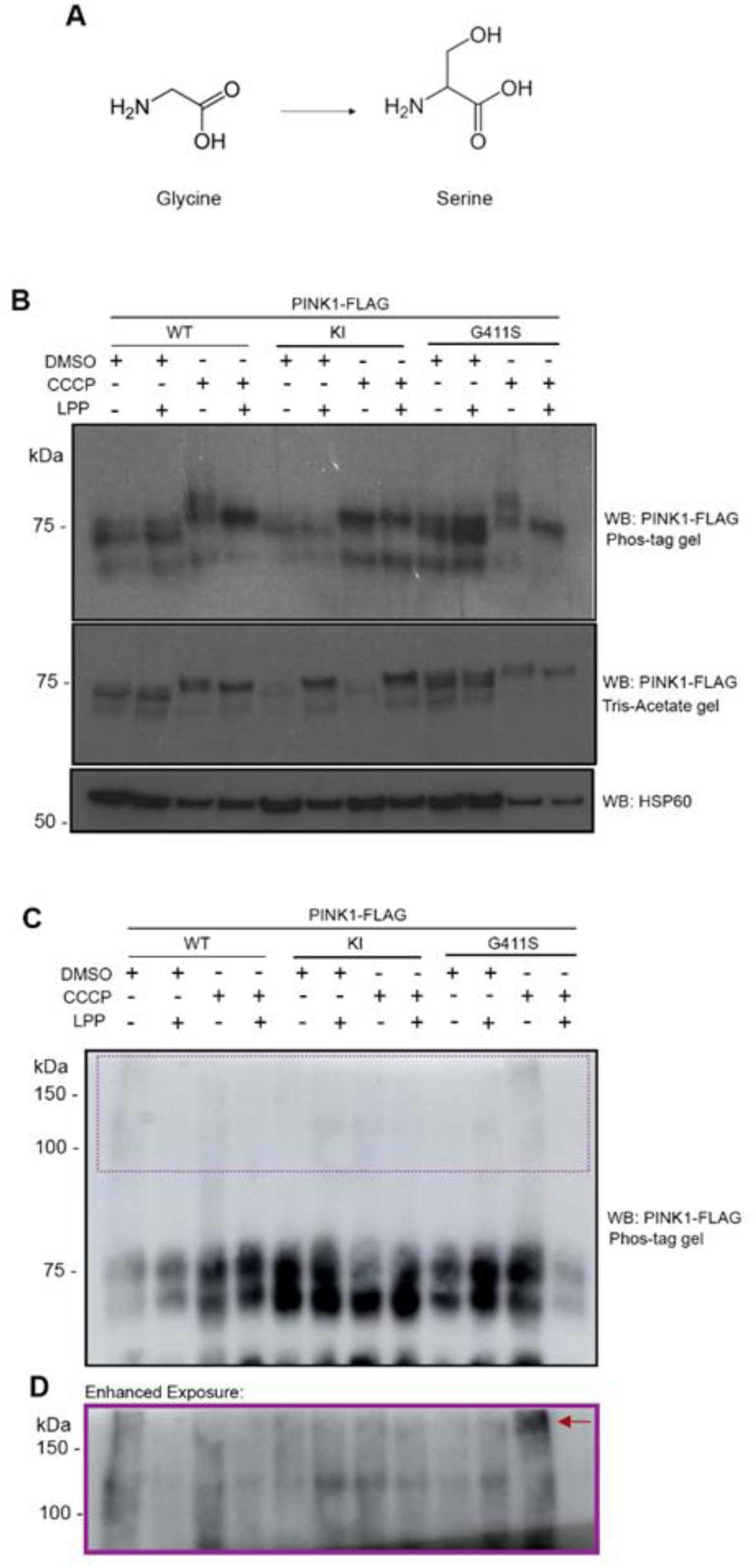
PINK1-G411S mutation generates an additonal phosphorylation site. A) Aminoacid change present in PINK1-G411S mutation, where in the residue 411 there is a replacement of a glycine for a serine. B) Cell lysates obtained from HEK293T cells transfected with PINK1-WT-FLAG, -KI-FLAG and -G411S-FLAG and incubated with 10µM of CCCP or DMSO, for 3hours, were treated with LPP and analyzed by SDS-PAGE on 7.5% Tris-acetate or 7,5% Phos-Tag gel and further immunoblotted for PINK1 detection using FLAG M2 antibody. Immunoblotting for mitochondrial marker HSP60 serves as protein loading control. C) Imunnoprecipitated samples obtained from HEK transfected with PINK1-WT-FLAG, -KI-FLAG and -G411S-FLAG and incubated with 10µM of CCCP or DMSO, for 3hours, were analyzed by SDS-PAGE on 7,5%Phos-Tag gel and further immunoblotted for PINK1 detection using FLAG M2 antibody. D) Enhanced exposure to detect higher molecular weight protei bands.

To test this hypothesis, we expressed PINK1-WT, -KI and -G411S with a C-terminal FLAG in HEK293T cells and analyzed its expression pattern on a Phos-Tag gel where negatively charged phosphorylated proteins migrate slower in comparison to their unphosphorylated counterpart. The altered migration pattern on the Phos-Tag gel and the clear decrease in intensity of the phosphorylated versus the non-phosphorylated PINK1 upon LPP and CCCP treatment suggest that, despite the detection of residual LPP-insensitive PINK1, the higher molecular weight band represents phosphorylated forms of PINK1 (Figure 6B). In PINK1-KI, phosphorylated forms of PINK1 are not detected, indicating that the observed phosphorylation event is dependent on PINK1 kinase activity (Figure 6B). To improve the detection of PINK1-G411S altered pattern, a FLAG immunoprecipitation was performed (Figure 6C), and LPP treated samples confirmed the presence of an additional high molecular weight band in PINK1-G411S that is absent in the PINK1-WT, indicating the possible presence of an additional phosphorylation site (Figure 6D).

### The additional phosphorylation site at the mutated G411S induces a stabilization of the ATP binding pocket

The putative PINK1 ATP-binding pocket includes 20 amino acids (I162, G163, K164, G165, C166, A168, V170, A217, K219, I275, M318, K319, N320, Y321, D366, N367, I368, L369, A383 and D384), as determined from sequence and structure predictions using the ATP bind algorithm(48) (Figure 7A,B). The two regulatory serine residues (Ser228 and Ser402)(21), and the glycine residue that is mutated to serine (G411) are located in the external boundaries of the ATP-binding pocket (Figure 7A). We hypothesize about the perturbations induced in the ATP binding pocket by the presence of the mutated and/or phosphorylated serine residues. Short-range atomistic molecular dynamics simulations were employed to determine the dynamics and stability of the ATP-binding pocket (Figure 7; Figure S6, S7, S8, S9, S10). The results, depicted in Figure 7, showed that in absence of phosphorylation, the ATP-binding cavity of the G411S mutant had an increased stability compared with the wild-type enzyme and demonstrated by the presence of a positive inter-residue correlation pattern among the involved aminoacids (Figure 7C,E), and a decrease in the number of aminoacids with low interaction times along the simulation (Figure 7C,E). This pattern is more dramatic when Ser411 is phosphorylated, and especially evident in the analysis of the Pearson inter-residue correlation (Figure 7E,F). Moreover, if we consider the tri-phosphorylated species of the PINK1 G411S mutant, the interaction time of the residues within the ATP-binding pocket was also incremented, when compared with the di-phosphorylated wild-type protein and with the native unphosphorylated (Figure 7C,D,G). All these structural changes are compatible with an increment of pocket rigidity and stability, directly related with the degree of phosphorylation of the PINK1 kinase.

**Figure 7.**
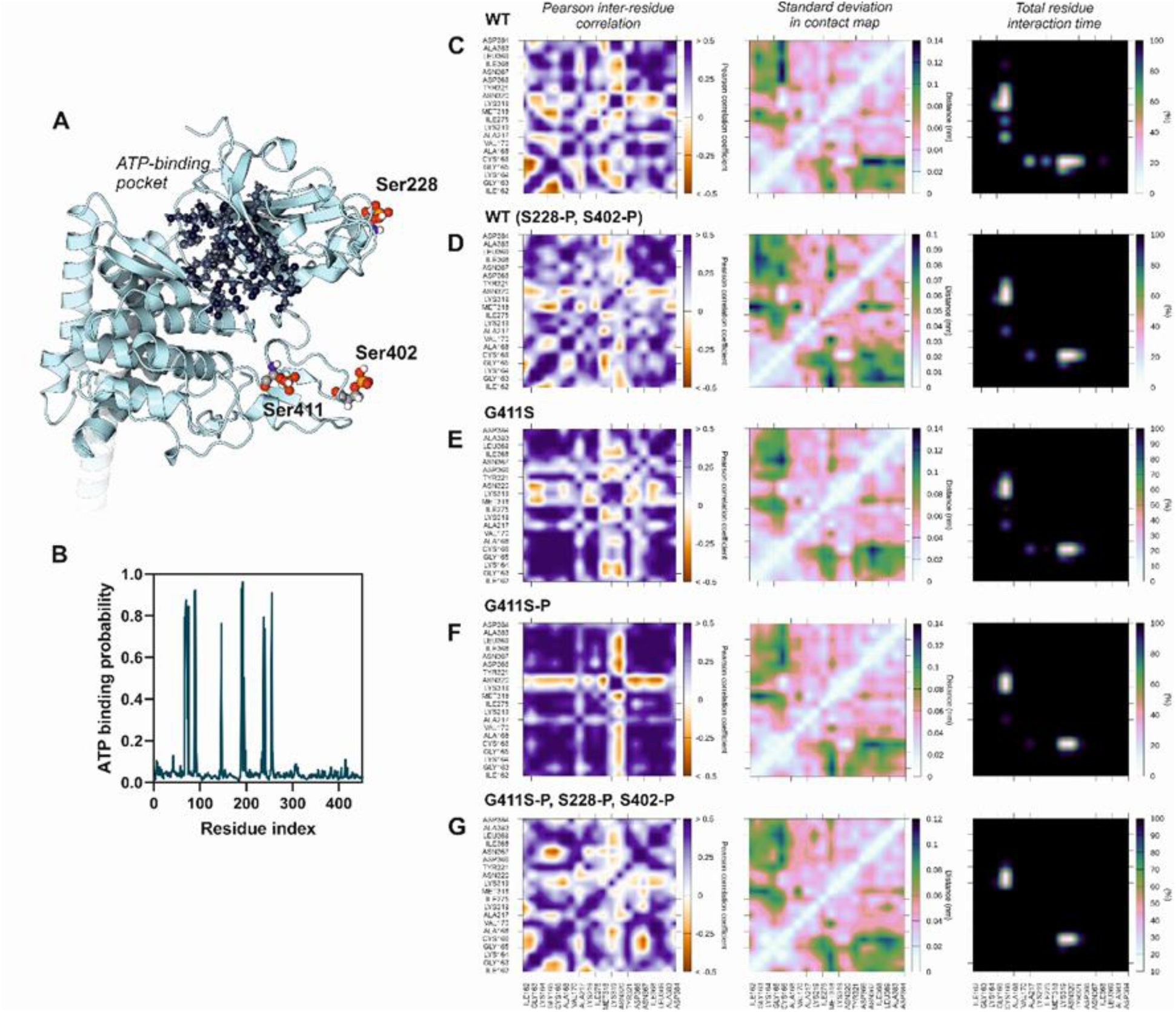
Short range molecular dynamics simulations of the ATP-binding pocket of the full atom model of PINK1. A) Structure model of PINK1 protein derived from AlphaFold predictions, showing the position of the three studied Serine residues (atom-colored sphere model) and the ATP-binding pocket (grey spheres); (B) ATP-binding probabilities along the PINK1 sequence, calculated with the ATPbind algorithm; trajectories analysis of the molecular dynamics simulations within the ATP-binding pocket, depicting the Pearson inter-residue correlations, the standard deviation in the contact map and the total residue interaction time along the simulation: (C) wild-type PINK1, (D) wild-type PINK with phosphorylated Ser228 and Ser402, (E) G411S mutant, (F) phosphorylated G411S mutant and (F) tri-phosphorylated mutated PINK1 at residues S228, S402 and S411.

## DISCUSSION

Parkinson’s Disease has a complex etiology, where both genetic and environmental factors play a role. Moreover, mitochondria dysfunctions and gene mutations, implicated in mitochondrial quality control (MQC), have been correlated with etiopathology and progression of PD, being one of them PINK1. The link between PINK1 and PD has been studied for several years, since the first PINK1 mutations was reported(3). These several years of research enlightened us about role of PINK1 in mitochondria and PD: *Drosophila* pink1 mutants show reduced life span, degeneration of flight muscles as well as dopaminergic neurons(17,18), while PINK1 KO mice have decreased dopamine levels and motor deficits(49) as well as impaired synaptic plasticity(50) and mitochondrial respiration(41,49). Several PINK1 mutations have been described as causative of autosomal-recessive PD, and even though some are characterized as compromising kinase activity or interfering with protein stability(6), there is still no clear mechanism of how it happens and if it is common for different mutations. Here we analyzed several mitochondria-PINK1 pathways to understand concretely how PINK1 mutations are disrupting mitochondria. Through an in vitro kinase assay we accessed PINK1 kinase activity in 8 human different mutations, and understood that majority of mutations, even outside the kinase domain, disrupt PINK1 ability to phosphorylate itself (Figure 1B,D) or its substrate Parkin (Figure 1C,E). Interestingly, PINK1-G411S mutation, localized in the activation loop, restores PINK1 kinase activity towards both substrates. PINK1 is one key player of MQC, so we further analyzed how this pathway may be disrupted with the presence of PD-related PINK1 mutations. For this, we investigated Parkin recruitment to mitochondria, as well as LC3 II receptor expression. Combining both results we observed, once again, that the mutant PINK1-G411S was able to recruit Parkin to mitochondria (Figure 2C), as well as LC3 II (Figure 2E). Proving, contrary to the other studied mutations, that PINK1-G411S was not disrupting MQC.

Taking this into consideration, we decided to pursue our studies analyzing only three different mutations: PINK1-C92F, because as previously described it is the only mutation outside the kinase domain; PINK1-G309D, because this mutation has previously shown to compromise kinase activity, behaving as a Kinase Inactive control(32); and the PINK1-G411S which does not disrupt MQC.

We further explored the role of PINK1 in mitochondrial morphology, where we saw a clear disruption of mitochondrial network (Figure 3B) and mass (Figure 3C) in MEF cells PINK1^-/-^ as well as in PINK1-G309D. Nevertheless, there were other studies reporting absence of mitochondrial morphological alterations in a PINK1^-/-^ background(40,41). One possible explanation for the mitochondrial network differences observed in our studies using MEF PINK1^-/-^ cells stably expression the PINK1 mutant with other reports could be either a due to the specific species used or variations in the degree of PINK1 inhibition, when using RNAi(40,41,45). Since PINK1 is responsible for phosphorylation of the Complex I subunit NDUFA10(51) and other more general deficiencies of the mitochondrial respiratory chain(41), we decided to analyze OXPHOS activity and expression in our PINK1 mutants of interest (Figure 4). We confirmed the results related to a reduced Complex I (NADH:ubiquinone oxidoreductase) enzymatic activity in the absence of PINK1(40,51) (Figure 4A). This decrease in Complex I activity is correlated with increased ROS production(52), as well as reduction in mitochondrial electrochemical gradient. Whereas Complex V seems only to be affected with every PINK1 mutations, indicating a compromised ATPase reductase activity in these cell lines (Figure 4B). Since PINK1 absence promotes a deficiency in the mitochondrial membrane potential (Δψm)(40), this metabolic switch to ATPase reductase may be a compensatory effort to preserve the Δψm(53). Nevertheless, when looking to OXPHOS protein expression levels (Figure 4C-E), no significant differences were reported for either Complex. However, we did observe a clear difference in NDUFA10 and NDFS6 (Complex I subunits) (Figure S4), which we postulate could be due to a misassembled Complex I. Dysfunctional complex in the dopaminergic neurons of the substantia nigra has been a hallmark of Parkinson’s disease(54). Recently, this dysfunction, was discovered as causative of progressive parkinsonism in mice with a conditional knockout in dopaminergic neurons for ndufs2, an essential subunit of Complex I(55). Moreover, PD patients’ samples revealed oxidatively damaged Complex I subunits, leading to a functionally impaired and misassembled Complex I(47). Being the ATPase reductase enzymatic activity disrupted in PINK1 mutations, we accessed ATP levels in these cells (Figure 4F). A clear impairment in PINK1^-/-^ MEF cells was observed, as an expected increase for PINK1-G309D and - G411S, meaning that this ATP accumulation could be due to lack of Complex V activity (Figure 4B). To test the effect of these PINK1 mutant forms on mitochondrial bioenergetics, we determined the mitochondrial respiration rates in response to bioenergetic modulators (Figure 5A), and no major differences were observed. PINK1^+/+^ are not producing (Figure 5D) or consuming (Figure 4B) more ATP, but still have higher ATP content than PINK1^-/-^, this may suggest that they have a higher balance of ATP:ADP ratio, which was previously observed in PINK1^-/-^ cortical neurons(56). The MEF PINK1^-/-^ have a higher spare respiration capacity (SRC) when compared to the PINK1^+/+^ counterpart (Figure 5F). Since SRC levels correlate with the degree of mitochondrial plasticity allowing bioenergetic adaptability in response to pathophysiological stress conditions (Figure S6), we hypothesize that MEF PINK1^-/-^ have adopted some compensatory mechanism to deal with the absence of PINK1. In fact, the loss of PINK1 impairs activity of mitochondrial respiration, which results in a decrease of ATP levels (Figure 4F) and increased ROS generation(57). This will impact glucose transporter and consequently substrate delivery to complexes I and II of ETC(40). To compensate and maintain Δψm ATP synthase is forced to consume rather than produce ATP (Figure 5B). Nevertheless, it would be interesting to confirm in high energy demanding tissues, such as brain, heart, and skeletal muscle, of aged PINK1^-/-^ animals or PINK1-PD samples, if there is an age-related decline in OXPHOS and consequent decrease in SRC, sensitizing the cells to surge in ATP demand, and increasing the risk of pathologies development. For the PINK1-G411S mutation, and considering the type of aminoacid change, we hypothesized that this PINK1 mutation could induce an additional phosphorylation site in PINK1. We tested this hypothesis using Phos-Tag gels (Figure 6B-D), and PINK1 molecular dynamics (Figure 7), and we were able to show, for the first time, that PINK1-G411S adds a phosphorylation site to PINK1. Interestingly, G2019S, one of the most common mutations in LRRK2, one important genetic risk factor for PD, results also in altered phosphorylation(58), and has an incomplete penetrance. The published data for PINK1-G411S is a bit controversial, some even consider this variant benign(25). Important to note that the G411S mutation in PINK1 was only identified as heterozygous polymorphisms of PD(59), which can either suggest this mutation is particularly damaging, or it may be considered a natural variant and not a disease-causing mutation(38). These variants could lead to a substantial increase in disease risk, without being pathogenic in all carriers(60) or clinical symptoms may be so subtle that they do not attracted medical attention(61). We hypothesized that this could be the case for PINK1-G411S, with this mutation having a dominant negative effect. During the preparation of this manuscript, Fiesel(62) and co-workers reported that PINK1-G411 site enhances enzymatic activity across different cell types, contradictory to previous data published by the same lab(24). The use of PINK1-G411S human fibroblast containing the additional phosphorylation site that we report support our theory of importance and presence of this mutation in PD-patient. Nevertheless, using cell based assays and structural computational analysis we additionally confirmed that PINK1-G411S mutation is responsible for altered stabilization and phosphorylation, of both PINK1 and Parkin. At the molecular level, the presence of an additional phosphorylation site in the G411S mutant, allowed us to determine an induced perturbation in the ATP-binding site that resulted in an increased rigidity and stability of the molecular pocket. The phosphorylation-dependent stabilization of ATP-binding pocket in kinases has been previously demonstrated in other human kinases, such as c-Src, supporting our findings(63). In sum, we were able to demonstrate that a putative stabilization of the ATP-binding pocket in the PINK1-G411S mutant increases the kinase domain’s rigidity leading to a fold-increase in PINK1’s overall enzymatic activity.

Our combined results provide a molecular framework to understand how amino acid changes in the backbone of PINK1, in particular for the G411 residue, has implications in the overall kinase activity. Thus, besides insights into the molecular mechanism of kinase regulation of PINK1, our findings open up new avenues for understanding the impact of PINK1 mutant forms and their implication in PD pathologies.

Concluding, different residues mutated in PINK1 can impair different mitochondrial pathways, as depending on the mutated residue a different crosstalk between PINK1 and its already described substrates will be deteriorated. In a disease context, our work strengthened the notion that each PD patient is a particular case and supports a future personalized medicine approach for these patients.

## EXPERIMENTAL PROCEDURE

### Cell Culture and Cell Lines

The HeLa-CrispR/Cas9-PINK1 cell line (here within referred to as HeLa PINK1^-/-^) were previously described(21). HeLa wild-type (WT), HeLa PINK1^-/-^, COS-1, HEK 293T and MEF cells were cultured at 37°C with 5% CO2 in DMEM/F12 medium (Thermo Fisher Scientific – Life technologies #LTI 11039-047) containing 10% fetal bovine serum (Gibco #A3840402). Stable cell lines expressing human PINK1-WT, PINK1-KI or mutant forms, PINK1-C92F and PINK1-G309D were generated by retroviral transduction of immortalized MEF cells derived from PINK1^-/-^ mice, previously described(40), and kindly provided by De Strooper Lab (VIB, Leuven). Stable expression of PINK1-G411S in MEF cell line were generated by retroviral transduction of immortalized MEF cells derived from PINK1^-/-^ mice as previously described(40). Selection of transduced cells was performed based on the resistance acquired to 5μM/mL of puromycin (Sigma-Aldrich #P8833). All cell lines were ideally manipulated with approximately 80% confluence. To induce mitochondrial membrane depolarization MEF and HeLa cells were treated with 25µM or 10µM CCCP (Sigma-Aldrich #C2759), respectively, or corresponding amount of DMSO (Sigma-Aldrich #41640) for control, 1h and 3h.

### Plasmids

The plasmids pcDNA3.1-hPINK1-FL-3xFLAG/Strep and a truncated form (ΔN) lacking first 113 aminoacids, pcDNA3.1-hPINK1-ΔN-3xFLAG/Strep; which one harbouring human PINK1 Wild Type (PINK1-WT) and human PINK1 Kinase Inactive (PINK1-KI were previously described(21)). To obtain kinase-inactive (KI) PINK1, lysine 219, in the ATP binding pocket, and aspartic acid 362, in the catalytic core, were both mutated to alanine (K219A/D362A)(21). PINK1-WT and PINK1-KI were also cloned into pMSCV retroviral vector(21). Mutant human PINK1 constructs were generated by QuickChange XL site-directed mutagenesis according to the manufacturer’s guidelines (Agilent Technologies #HPA 200516). The following constructs were made: pMSCVpuro-hPINK1, pcDNA3.1-hPINK1-FL-3xFLAG/Strep or pcDNA3.1-hPINK1-ΔN-3xFLAG/Strep using primers described in Supplemental Table 1. After PCR amplification, the parental non-mutated DNA is specifically digested, and the remaining mutated plasmid DNA is transformed. Subsequently, the DNA of single colonies is analyzed for the presence of the mutation via restriction digest where possible. The Ubiquitin-like binding domain (Ubl) of Parkin was obtained by PCR amplification of amino acids 1–108 from the pCMV-Parkin plasmid (Origene). The PCR product was further cloned into pGEX-4T-1 (Addgene) in-frame with the N-terminal GST-tagged fusion construct(21). All plasmids were confirmed by performing sequencing analysis. Mitochondrial targeted DsRed (mitoDsRed) plasmid was gift from L. Scorrano Lab (Venetian Institute of Molecular Medicine, Pádua, Italy) Plasmids were transiently transfected using TransIT transfection reagent (Mirus Bio #MIR 2300) according to the instructions of the manufacturer. Briefly, 1µg of DNA plasmid per 3µl of transfection reagent ratio was used.

### Cell Lysates

MEF cells, HEK293T and HeLa cell lines were harvested with 100-150μL Lysis Buffer (50mM Tris-HCl pH 7.4, 5mM EDTA, 150mM NaCl, 1% Triton X-100, Protease inhibitors (1/100)) and pelleted for 3min, 800xg at 4°C. After 1h of incubation on ice, lysates were centrifuged at 10,000xg for 10min at 4°C. Supernatants were collected and protein concentration was accessed using Bradford Protein Assay (Bio-Rad #5000007). Samples were stored at -20°C.

### Immunoprecipitation

Cell lysates obtained as previous described, from HEK293T cells transfected with PINK1-WT-FLAG, KI-FLAG and G411S-FLAG, were incubated with FLAG beads (Sigma-Aldrich #M8823) with end-over-end rotation, overnight at 4°C. Samples were used for dephosphorylation.

### Protein Dephosphorylation

The cell lysates or purified PINK1 bound to FLAG beads were incubated at 30°C for 1h with 200 units of ʎ protein phosphatase (LPP) in a total reaction of 100µL following the instructions of the manufacturer (New England Biolabs #P0753).

### Subcellular Fractionation

MEF PINK1-WT and mutant forms were harvested with Isolation buffer (IB) (0.2M sucrose, 10mM Tris/Mops pH=7.4, 0.5mM EGTA/Tris pH=7.4) and centrifuged for 10min, 600xg at 4°C. Pelleted cells were resuspended in IB and homogenized using a Teflon pestle with 30-40 strokes at 1000rpm. Homogenates were centrifuged at 600xg for 10min at 4°C. Supernatants were centrifuged at 7000xg for 10min at 4°C. Pellets was resuspended in IB and washed at 7000xg for 10min at 4°C (Adapted from(64)). Pelleted mitochondria were resuspended in IB.

### SDS-PAGE and Immunoblotting on Tris Acetate and Phos-Tag Gels

Equal protein amounts of cell lysates and mitochondria-enriched fractions were separated using SDS-PAGE gels (PreCast Mini-PROTEAN Tris Glycine TGX, 4-15% (Bio-Rad #4561083); PreCast Mini-PROTEAN Tris-Glycine TGX 7,5% (Bio-Rad #4561026) and PreCast Bolt 10% Bis-Tris (Invitrogen #NW00105BOX) percentages previously described) and transferred to Nitrocellulose (GE Healthcare #10600001) or PVDF membranes (GE Healthcare # 10600023). Blocking was performed for 1h using 5% dried milk or Bovine Serum Albumin (Sigma-Aldrich #A9647) in TBS-T (50mM Tris-HCl pH 7.5; 150mM NaCl, 0.1% Tween-20 (Sigma-Aldrich #P1379)) and consequently membranes were incubated with primary antibodies (described in Supplemental Table 2) overnight at 4°C in blocking solution. Next, membranes were incubated with species-compatible secondary HRP-couples antibodies for 2h at room temperature (RT).

For analysis on PhosTag gels, samples were incubated for 10min at 70°C in Tris/glycine SDS samples buffer (BioRad) with 4% β-mercaptoethanol. Resolution gels with 7.5% acrylamide (Acryl/Bis 29:1; BioRad #A3574), 50µM Phos-TagTM AAL-107 (Wako Chemicals #300-93523) and 100µM MnCl2 were casted according to the instructions of the manufacturer, including 4.5% stacking gels. Mn2+-Phos-TagTM interacts with phosphorylated proteins, altering the pattern of migration. Prior to transfer to a PVDF membrane, gels were washed for 10min with gentle agitation in transfer buffer containing 1mM of EDTA to eliminate manganese.

Enhanced chemiluminescence method was used for protein detection (ECL - GE Healthcare #RPN2106), membranes were incubated with ECL mix for 1minute at room temperature and then, the protein bands were detected using automatic film processor - Curix 60 (AGFA) or AmershamTM ImageQuantTM 800 imager. Semi-quantification was performed using Image studio lite version 5.2.5 software.

### Immunofluorescence

HeLa and MEF cell lines cultured on glass coverslips were fixed for 10min using 4% paraformaldehyde (Electron Microscopy Sciences #15710), in PBS followed by permeabilization for 20minutes with 0,1% Triton X-100 (Sigma-Aldrich #T8787) in PBS and blocked for 1h with Blocking Buffer (1x PBS, 0.2% Tissue culture grade gelatine, 2% FBS, 2% BSA, 0.3% Triton X-100 with 5% Goat Serum (Dako #X090710)). Cells were incubated for 2h with primary antibody diluted in Blocking Buffer with 5% Goat Serum (Dako #X090710) (antibodies are dilutions used are listed in Table 2). Cells were washed 3x in PBS and further incubated for 2h with specie-specific Alexa Fluor secondary antibodies (Molecular Probes). Cells were imaged using a Zeiss LSM 710 confocal microscope and images were analyzed using ImageJ 1.53c software. Co-localization parameters were determined using JACop Plug-in(65) which evaluates Mander’s overlap coefficient, which is based on Pearson’s collection coefficient.

### Mitochondria live-imaging

For live imaging, cells were transfected with fluorescent mitochondrially targeted MitoDsRed plasmid as described above. Images were acquired on a Zeiss Cell Observer SD and analyzed using ImageJ 1.53c software, where an automatic threshold function was used. Mitochondria mass was quantified by dividing the number of pixels corresponding to mitochondrial staining by the total area of the cell.

### Morphometric analysis

Morphology analysis was performed as previously described in Morais et al, 2009 (40), (66) and examined in LSM710 confocal microscope.

### Antibodies

Antibodies were used for Western Blot analysis (WB) and Immunofluorescence (IF) are described in the Supplemental Table 2.

### Parkin expression and purification

Recombinant expression and purification of Parkin was performed as previously described(21). Briefly, BL21 bacteria strain was transformed with the pGEX-4T-1 vector expressing an N-terminal GST-tagged Ubl-domain of Parkin. Induction was performed with 100μM IPTG, at 37°C with 280rpm of agitation for 2h. After centrifugation, bacterial pellets were lyzed in 50mM Tris-HCl pH 7.5, 150mM NaCl, 1% Triton X-100, 2mM EDTA, 0.1% β-mercaptoethanol, 0.2mM PMSF and 1mM benzamidine. GST-Ubl Parkin was purified using Glutathione Sepharose™ 4B (GE Healthcare #17-0756-01), according to manufacturer’s instructions. Control samples were retained in every purification step, quality, and purity were evaluated via western blot.

### Human PINK1 purification and in vitro kinase assay

This procedure was performed as previously described(21). Briefly, COS-1 cells were transfected with TransiT as previously described, and 48h post transfections, cells were washed and harvested in PBS. After a 10min centrifugation at 200xg, cell pellets were resuspended in Lysis buffer (25mM Tris-HCl pH7.5, 150mM NaCl, 5mM NaF, 1mM MgCl2, 1mM MnCl2, 0.5% Igepal-NP40 (Sigma), 50mg/L DNAse (Sigma), 50mg/L RNAse (Sigma), 1mM DTT), with 20% protease inhibitor cocktail for mammalian cell and tissue extraction (Sigma-Aldrich #P8340), 2x complete protease inhibitor (Sigma-Aldrich #4693132001), 4x PhosSTOP tablets (Roche #4906837001) and homogenized using a 22-G needle in 5 strokes. Lysates were centrifuged for 25min at 20,000g, and then incubated for 45min at 4°C with FLAG-magnetic beads (Sigma-Aldrich #M8823). The unbound fraction was discarded, and beads were washed 2x with Lysis buffer and 3 times with kinase assay buffer (50mM Tris-HCl pH7.5, 150mM NaCl, 10mM MgCl2, 3mM MnCl2 and 0.5mM DTT). The kinase assay was performed immediately after the binding step where purified hPINK1-FLAG bound to the beads was incubated with 2μg Parkin, 10mM DTT and 100μM ATP containing 5μCi [γ-32P] ATP and incubated for 1h at 22°C. Samples were analyzed by SDS-PAGE followed by Western blotting. Incorporation of radiolabelled phosphate was assessed via a storage phosphor screen and development using an Amersham Typhoon IP (GE Healthcare Life Sciences). Image studio lite version 5.2.5 software was used for signal quantification.

### Bioenergetic Measurements

Cellular bioenergetic measurements were performed using the Seahorse XFe24 Analyzer (Agilent Technologies), using the XF Cell Mito Stress Test. Briefly, MEF PINK1^-/-^, MEF PINK1^+/+^, MEF PINK1-WT, -KI and mutant cell lines were seeded in Seahorse XF cell culture microplates (Agilent Technologies #102340-100) at a density of 10,000 cells/well one day prior to the analysis. On the day of the assay, cell culture medium was replaced by Seahorse XF DMEM assay medium (Agilent Technologies # HPA102353100) supplemented with 10mM D-Glucose (Sigma-Aldrich #G7021), 1mM sodium pyruvate (Thermo Fisher Scientific #LTID 11360-039) and 2mM L-glutamine (Thermo Fisher Scientific #LTID 25030-024) and incubated in CO2-free incubator for 1h prior to assay. Baseline of oxygen consumption was determined, followed by sequential injections of 1μM oligomycin (Sigma-Aldrich #O4876-5MG), 2μM FCCP (Sigma-Aldrich #C2920) and 1μM rotenone (Sigma-Aldrich #R8875) + 1μM antimycin A (Sigma-Aldrich #A8674) to reveal the key parameters showing metabolic function. Each measurement was normalized to protein content. Afterwards, based on response to several injections multiple parameters were calculated. After the basal measurements, there is a 1st injection (Oligomycin), which impacts on electron flow through ETC, resulting in a decrease of OCR connected to cellular ATP production. The response to FCCP (2nd injection) is used to calculate maximum respiration, since flow through the ETC is uninhibited and oxygen consumption by Complex IV reaches the maximum. In response to FCCP it can also be calculated the spare respiratory capacity, defined as the difference between maximal respiration and basal respiration. The last injection (mixture of rotenone and antimycin A) allows to calculate the respiration driven from processes outside mitochondria, the nonmitochondrial respiration.

### ATP content

ATP content was performed as previously described(51) using the ATP Determination Kit (Molecular Probes #A22066). Briefly, cells were lysed with Extraction buffer (6M Guanidine-HCl, 100mM Tris-HCl pH7.8, 4mM EDTA) and protein content was accessed using BCA Protein Assay (Thermo Fisher Scientific #23225). Then 5μg of protein was incubated with 1X Reaction Buffer, 0.1M DTT, 0.5mL of 10mM D-luciferin and 2.5μL/mL of firefly luciferase. Luminescence was accessed using the SynergyHTX equipment.

### Enzymatic Activity

Citrate synthase (CS), NADH:ubiquinone oxidoreductase (complex I) and F1F0 ATP synthase (complex V) enzymatic activities were determined by colorimetric measurements as previously described(40). In brief, for citrate synthase the reaction mixture contained 50μg of mitochondria, 0,1M Tris-HCl pH8.1, 0.1% Triton X-100, 0.2mM 5,5′-Dithiobis(2-nitrobenzoic acid) and 0.15mM acetyl-CoA. After 2min stabilization at 37°C, citrate synthase reaction started by adding 10mM oxaloacetate (Sigma-Aldrich #O4126). Measurements were performed at 412nm for 8 min. Complex I activity was measured by monitoring NADH oxidation in the presence of decyluquinone at 340nm for 45 min. The reaction mix contained 50μg mitochondria, 0,1M K-phosphate buffer pH7.5, 2,5mg/mL BSA (Sigma-Aldrich #A6003), 5mM MgCl2 (Sigma-Aldrich #M9272), 100μM NADH (Roche #10128023001), 0,6mM KCN (VWR #26802.234) and 10μM Rotenone (Sigma-Aldrich #R8875). After 1min stabilization at 37C, 130mM Decylubiquinone (Sigma-Aldrich #D7911) was added. Complex I activity was calculated using the difference between the measured rates in the absence vs. presence of the Complex I inhibitor rotenone. Complex V activity was determined by using 50μg of mitochondria, 10mM Complex V buffer (0,1M Tris (Sigma-Aldrich #T1503), 0,01M MgCl2 (Sigma-Aldrich #M9272), 0,02M KCl (Sigma-Aldrich #P3911), 2mM PEP (Sigma-Aldrich #P7252), 0,2mM NADH (Roche #10128023001), 7,5U/mL PKinase (Sigma-Aldrich #L2500), 12μM oligomycin (Sigma-Aldrich #O4876), 1mM ATP (Sigma-Aldrich #A7699)). Activity was measured at 340 nm for 1h. All spectrophotometric measurements were performed at 37°C. Enzymatic activity was calculated by the difference between colorimetric measurements without and with inhibitors, for both analyzed complexes, and normalized to Citrate Synthase activity.

### Molecular Dynamics

Atomistic short-range molecular dynamics (MD) simulations were performed on the Alphafold-generated(67) human PINK1 model structure (Alphafold database ID: AF-Q9BXM7-F1). Segments predicted with confidence levels below 70 were removed from the model, comprising residues 1-95 and 180-213. Selected mutants were generated by aminoacid replacement within the structure using Coot software(68). Generation of topology and molecular dynamics parameters for the monomeric and dimeric ensembles was performed with CHARMM-GUI(69). The protein model was placed in the center of a cubic box, and the system solvated in TIP3 water molecules and 0.15 MKCl, imposing an edge distance of 10nm. CHARMM36 force field was used to obtain the simulation parameters for the protein and solvent. All amino acid residues were kept in their standard protonation states compatible with a physiological pH (e.g. all lysine, arginine, glutamate, and aspartate residues were charged). The model system size was roughly 183,000 atoms.

All the simulations were performed with GROMACS 2021(70). The model systems were first energy minimized during 1000 steps and the temperature was further equilibrated to 310 K in NVT ensemble using V-rescale thermostat during a 1 ns simulation. The systems were further equilibrated in NPT ensemble using V-rescale thermostat and Berendsen barostat for an additional 1,000 ps, followed by removal of all constraints and further NPT equilibration for 2,000 ps. In the production phase, no constraints were applied, and the temperature and pressure were kept at 310 K and 1 atm using the Nose-Hoover thermostat and Parrinello–Rahman barostat. The timestep of the simulations was 2 fs which was achieved using the LINCS algorithm(71). Production phase simulations were run for 10 ns in the monomeric system and for 20 ns in the dimeric ensemble. The particle-mesh Ewald method was used to handle electrostatic interactions with a 12 Å cutoff. The van der Waals cutoff was also 12 Å with switching distance at 10 Å. Generated simulation trajectories were analysed and visualized using GROMACS(70), CorrelationPlus(72) and CONAN softwares(73).

### Statistical analysis

Statistical significance was determined using GraphPad Prism7 software, through Student’s t-test or one way ANOVA with Dunnett’s multiple comparison test as indicated (* P < 0.05; ** P < 0.01; *** P < 0.001; **** P < 0.0001; n.s. – non significant). Data are shown as mean +/- standard errors of the mean (SEM), with 95% of confidence interval, in a minimum of 3 independent biological replicates.

## DATA AVAILABILITY STATEMENT

Data is contained within the article or Supplementary Material. Additionally, the datasets generated during and/or analysed during the current study are available from the corresponding author on reasonable request.

## SUPPORTING INFORMATION

This article contains supporting information.

## ACKNOWLEDGMENTS

We would like to thank Bart de Strooper, Liesbeth Aerts and Kathleen Creassearts (VIB-KULeuven, Belgium) for providing the PINK1 related expression plasmids and HeLa PINK1 null cells. We thank members of the VMorais Lab for fruitful discussions. We would like to thank the BioImaging Facility, with a special thanks to José Rino, António Temudo and Ana Nascimento, and we also acknowledge the funding PPBI-POCI-01-0145-FEDER-022122.

## AUTHOR CONTRIBUTIONS

Conceptualization, F.B.G and V.A.M.; Methodology, F.B.G and F.J.E.; Software, F.J.E.; Validation, F.B.G and F.J.E.; Formal Analysis, F.B.G and F.J.E.; Investigation, F.B.G and V.A.M.; Resources, F.B.G and V.A.M.; Data Curation, F.B.G, V.A.M. and F.J.E.; Writing – Original Draft Preparation, F.B.G and F.J.E.; Writing – Review & Editing, F.B.G, V.A.M. and F.J.E..; Supervision, V.A.M.; Project Administration, V.A.M.; Funding Acquisition, V.A.M. All authors have read and agreed to the published version of the manuscript.

## FUNDING AND ADDITIONAL INFORMATION

This research received financial support from Fundação para a Ciência e Tecnologia with the following Grant references: for FBG SFRH/BD/134316/2017, COVID/BD/152472/2022, and IMM/BI/1c-2023; for VAM IF/01693/2014; and Funding Grant reference PTDC/BIA-CEL/31230/2017 by Fundação para a Ciência e a Tecnologia (FCT). Funding was attained from the following research grants: ERC-StG-679168 by European Research Council and EMBO-IG/3309 by European Molecular Biology Organization.

