## Supplemental Information for "PINK1-G411S mutant increases kinase stability and enhances mitochondrial-linked functions"

**TITLE**

*****Correspondencing author:

Vanessa A. Morais,

### Figure S1

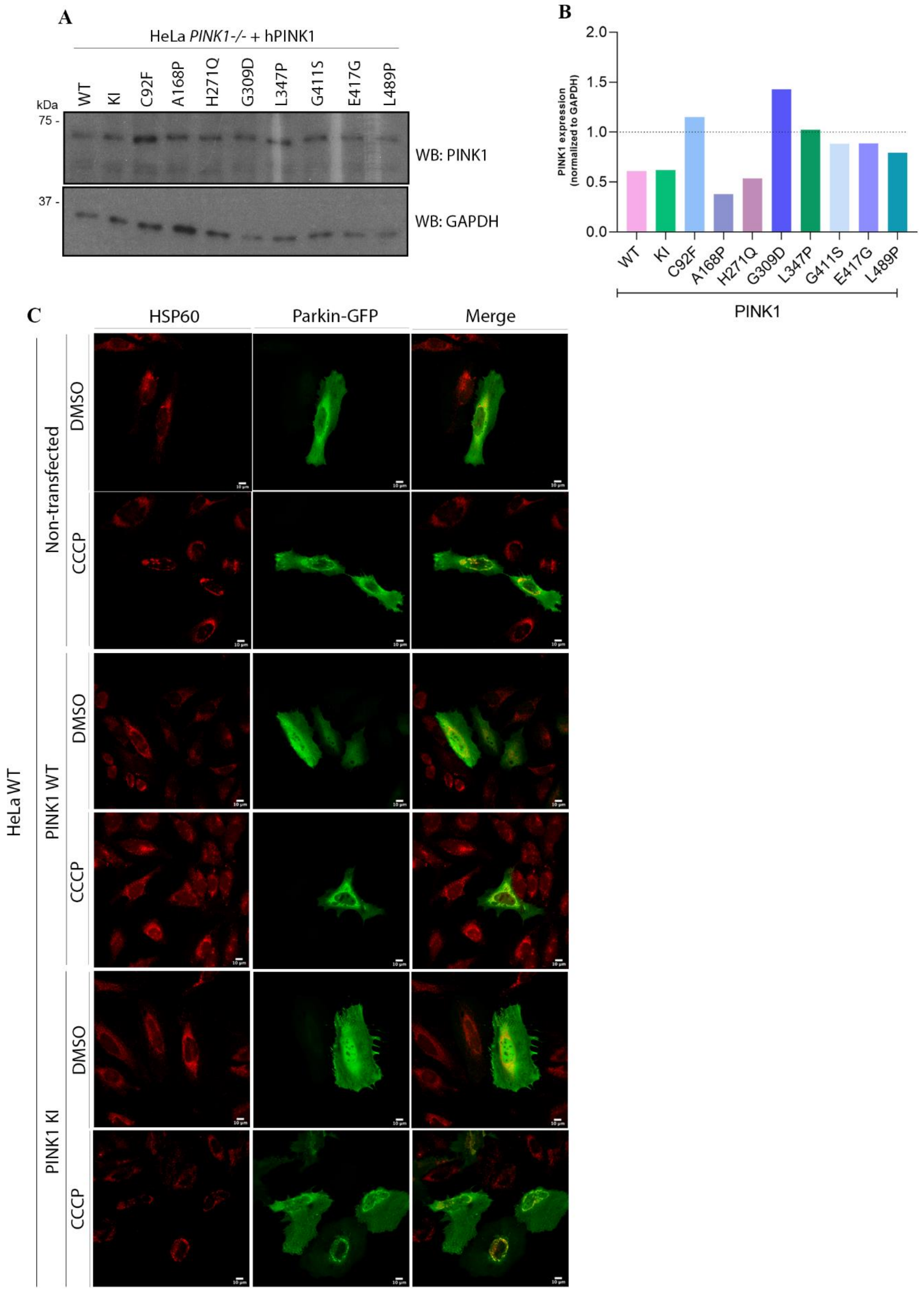
A
HeLa PINK1-/- + hPINK1
WT
KI
C92F
A168P
H271Q
G309D
L347P
G411S
E417G
L489P
kDa
75 -
37 -
WB: PINK1
WB: GAPDH
B
PINK1 expression (normalized to GAPDH)
2.0
1.5
1.0
0.5
0.0
PINK1
C
HSP60
Parkin-GFP
Merge
HeLa WT
Non-transfected
PINK1 WT
PINK1 KI
DMSO
CCCP

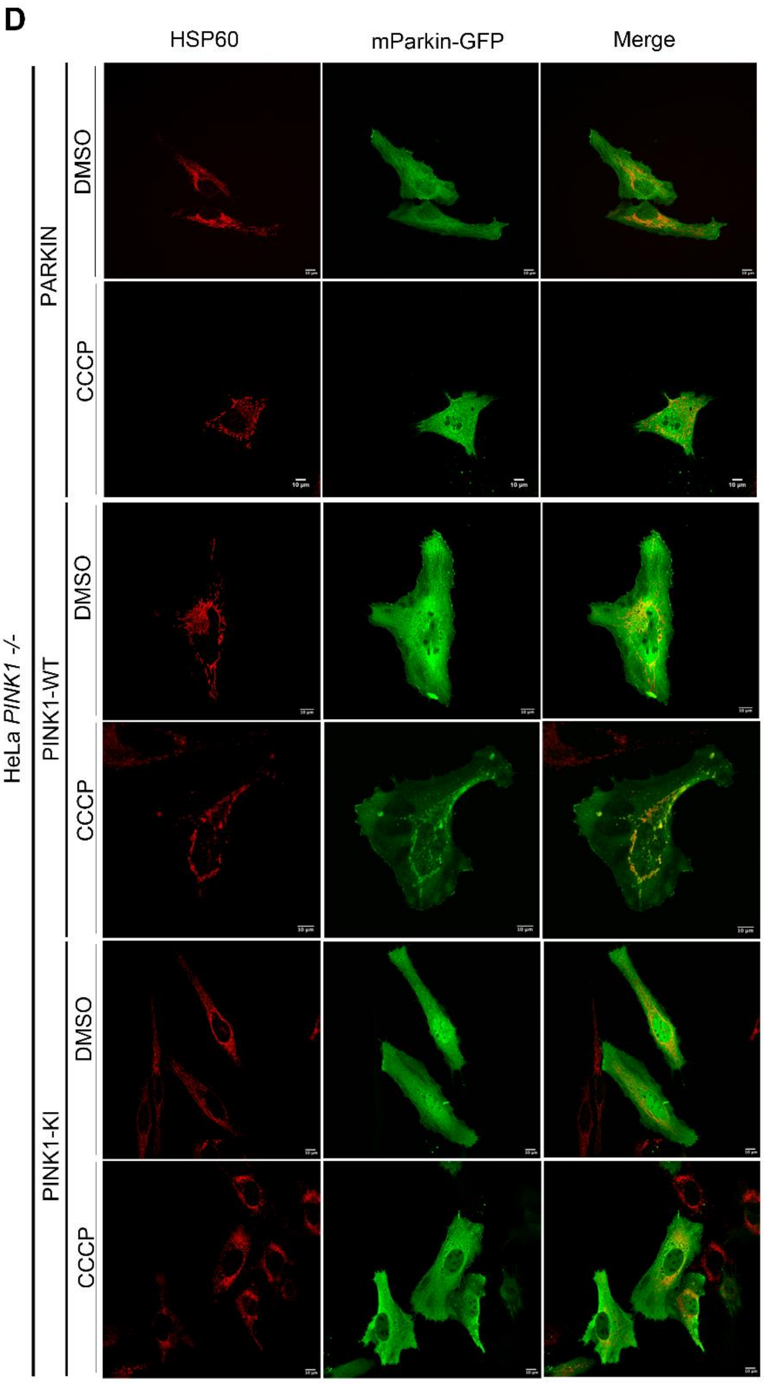
D
HSP60
mParkin-GFP
Merge
HeLa PINK1 -/-
PARKIN
PINK1-WT
PINK1-KI
DMSO
CCCP
DMSO
CCCP
DMSO
CCCP
10 µm

E

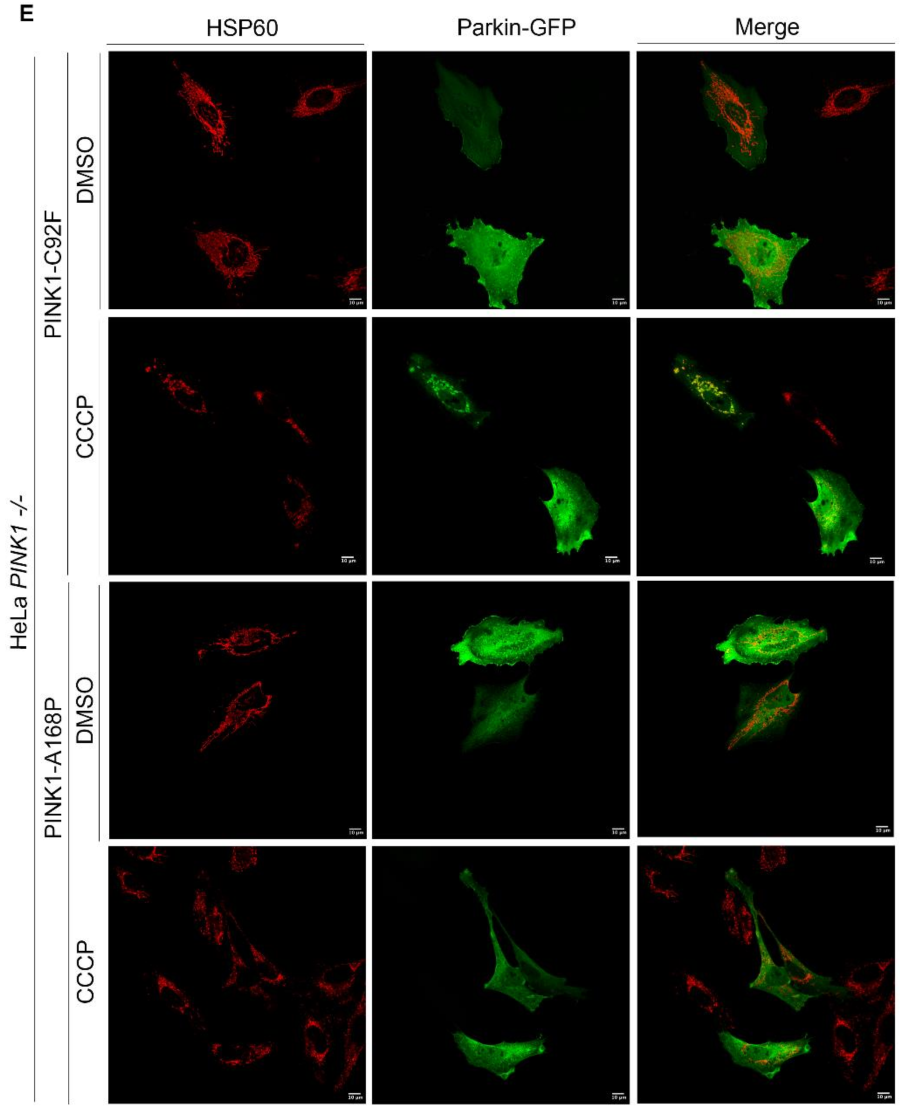
HSP60
Parkin-GFP
Merge
HeLa PINK1 -/-
PINK1-C92F
DMSO
CCCP
PINK1-A168P
DMSO
CCCP

F

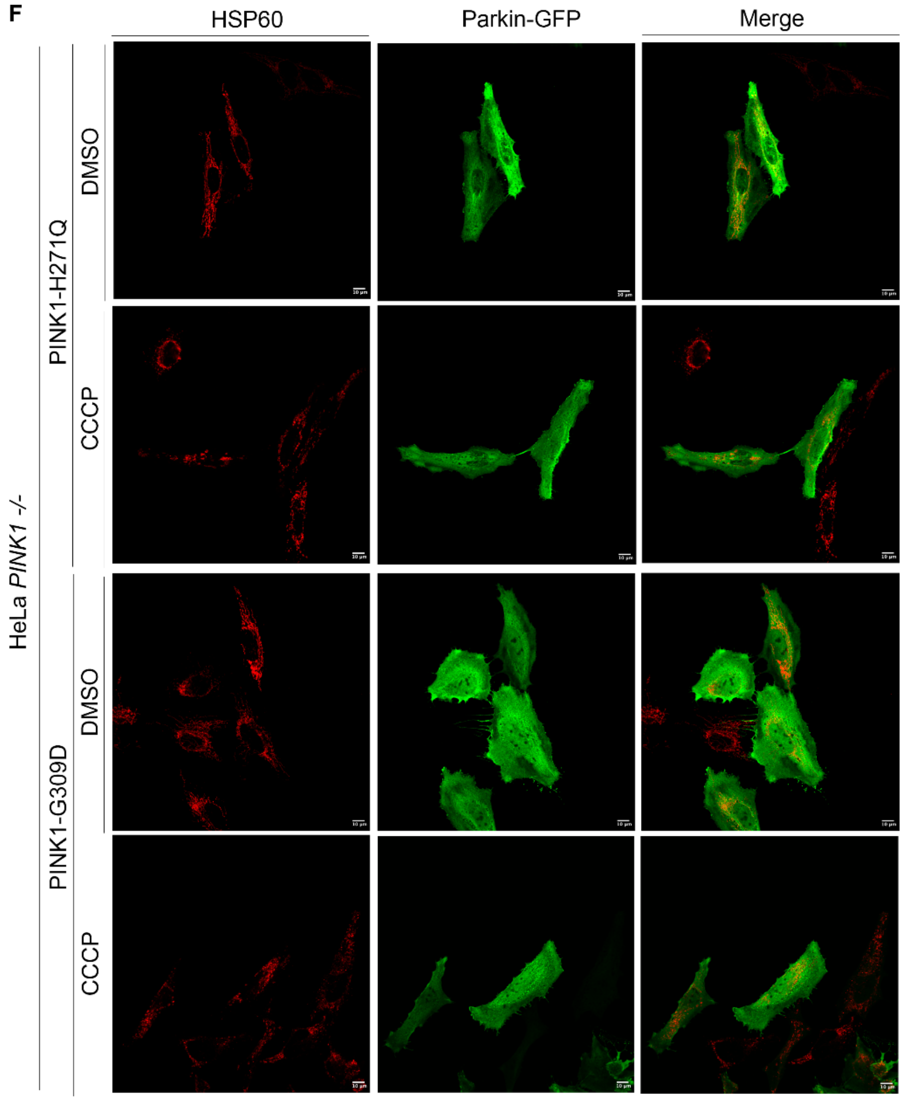

HSP60
Parkin-GFP
Merge
HeLa *PINK1 -/-*
PINK1-H271Q
DMSO
CCCP
PINK1-G309D
DMSO
CCCP

G

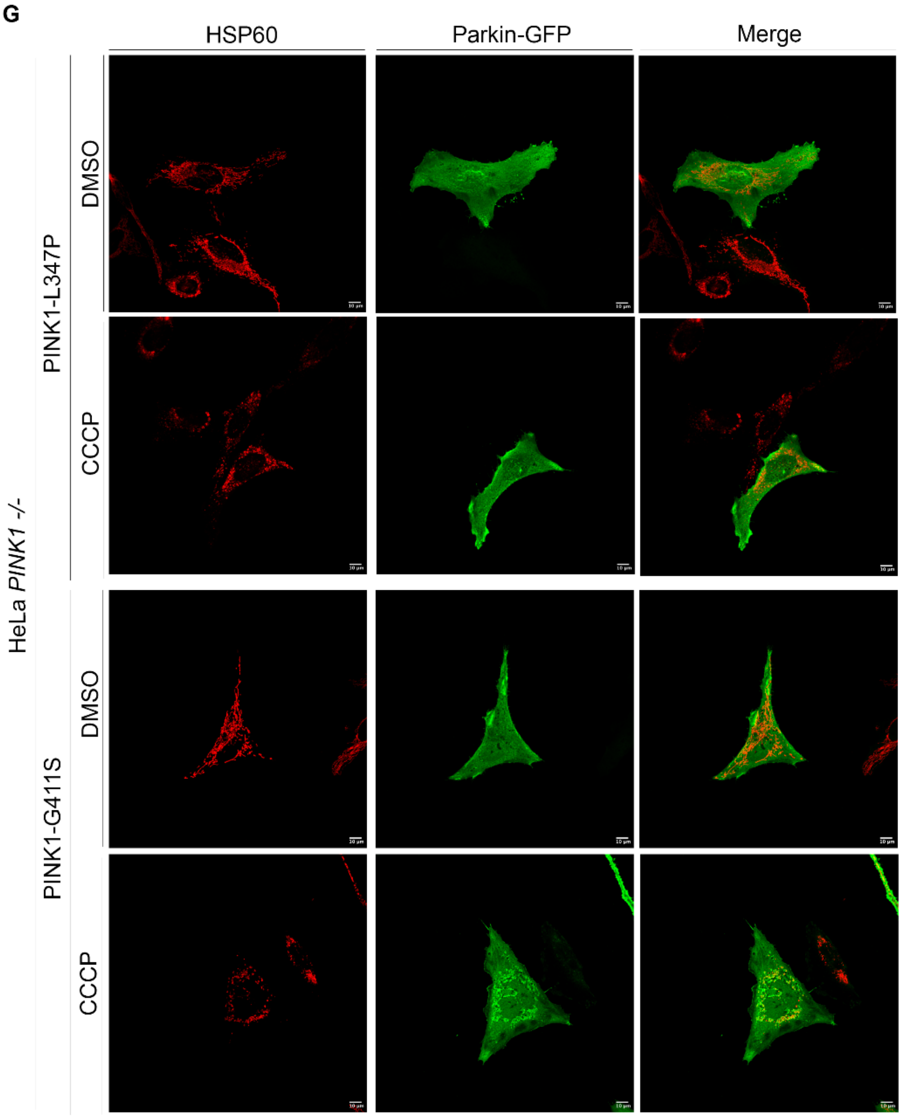

HSP60
Parkin-GFP
Merge
HeLa PINK1 -/-
PINK1-L347P
PINK1-G411S
DMSO
CCCP
DMSO
CCCP

H

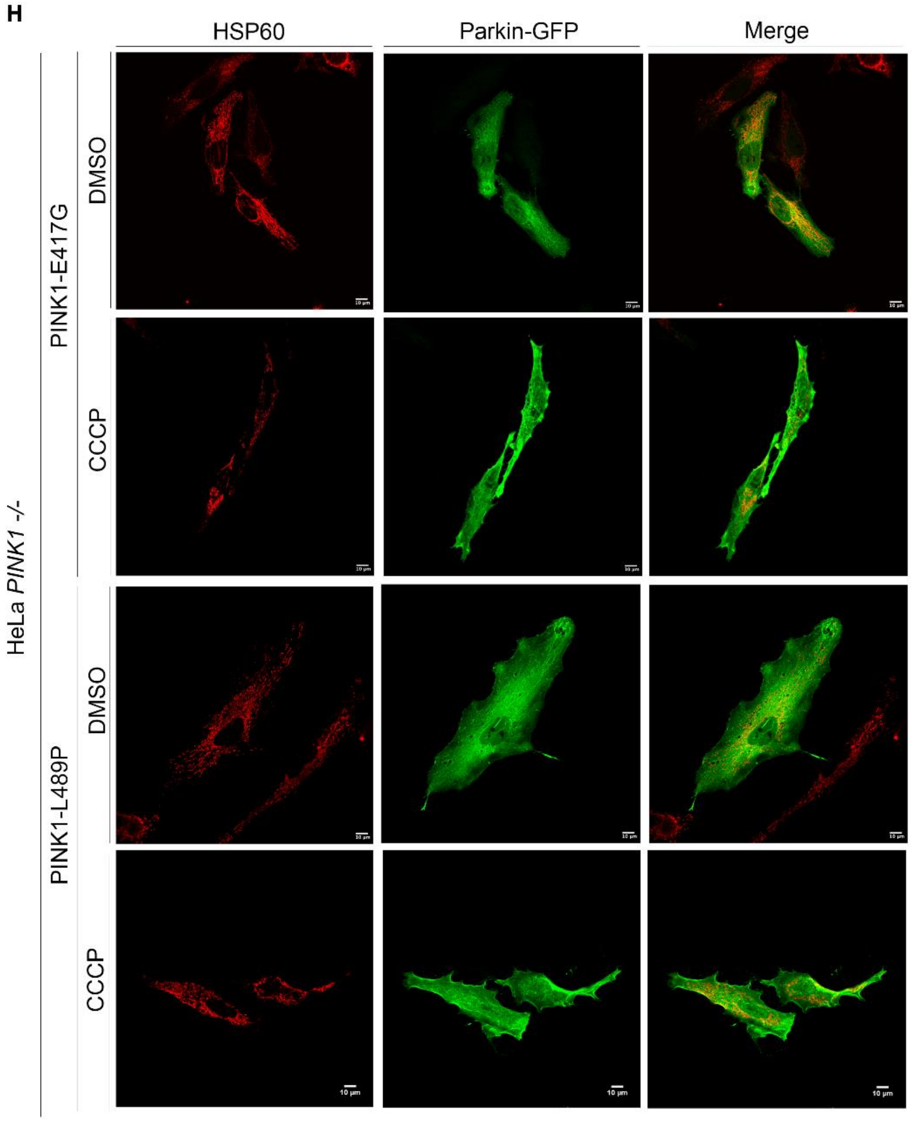


***Supplemental Figure S1*** – *Analysis of PINK1 protein levels.*

*A) HeLa PINK1$^{-/-}$ cells were transfected with pMSCV PINK1 WT, KI or studied PINK1clinical mutations, and were analyzed by SDS-PAGE on 7.5% Tris acetate gel and probbed for PINK1 and HSP60, as loading control marker. B) Quantification of PINK1 expression in HeLa PINK1$^{-/-}$ cells. C-H) HeLa WT or HeLa PINK1-/- cells were transfected with PINK1 WT, KI or PINK1 clinical mutants, and treated with DMSO or 10μM CCCP, to induce depolarization (Scale bar=10μm).*

### Figure S2

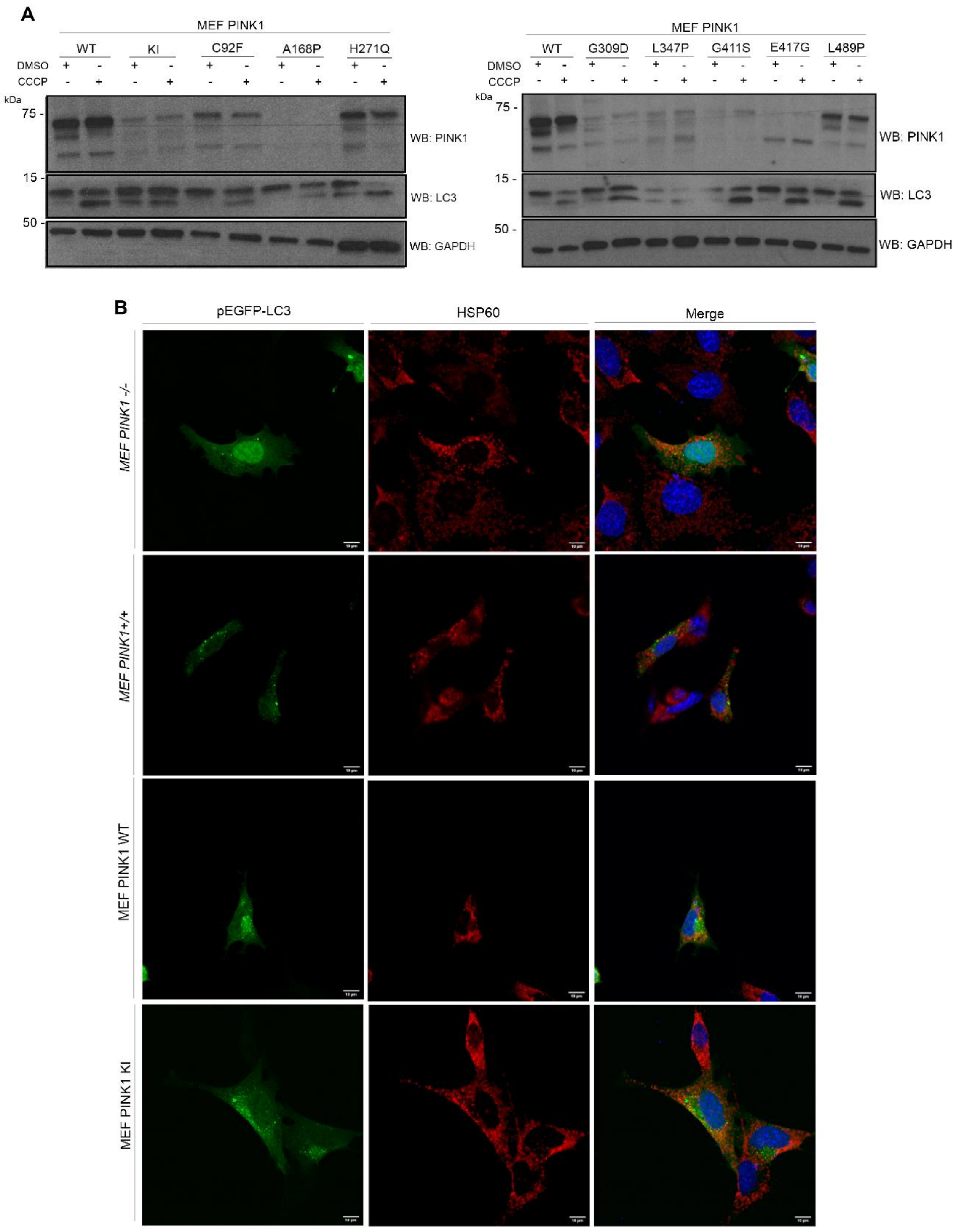
A
MEF PINK1
WT
KI
C92F
A168P
H271Q
DMSO
CCCP
kDa
75 -
15 -
50 -
WB: PINK1
WB: LC3
WB: GAPDH
MEF PINK1
WT
G309D
L347P
G411S
E417G
L489P
B
pEGFP-LC3
HSP60
Merge
MEF PINK1 -/-
MEF PINK1+/+
MEF PINK1 WT
MEF PINK1 KI

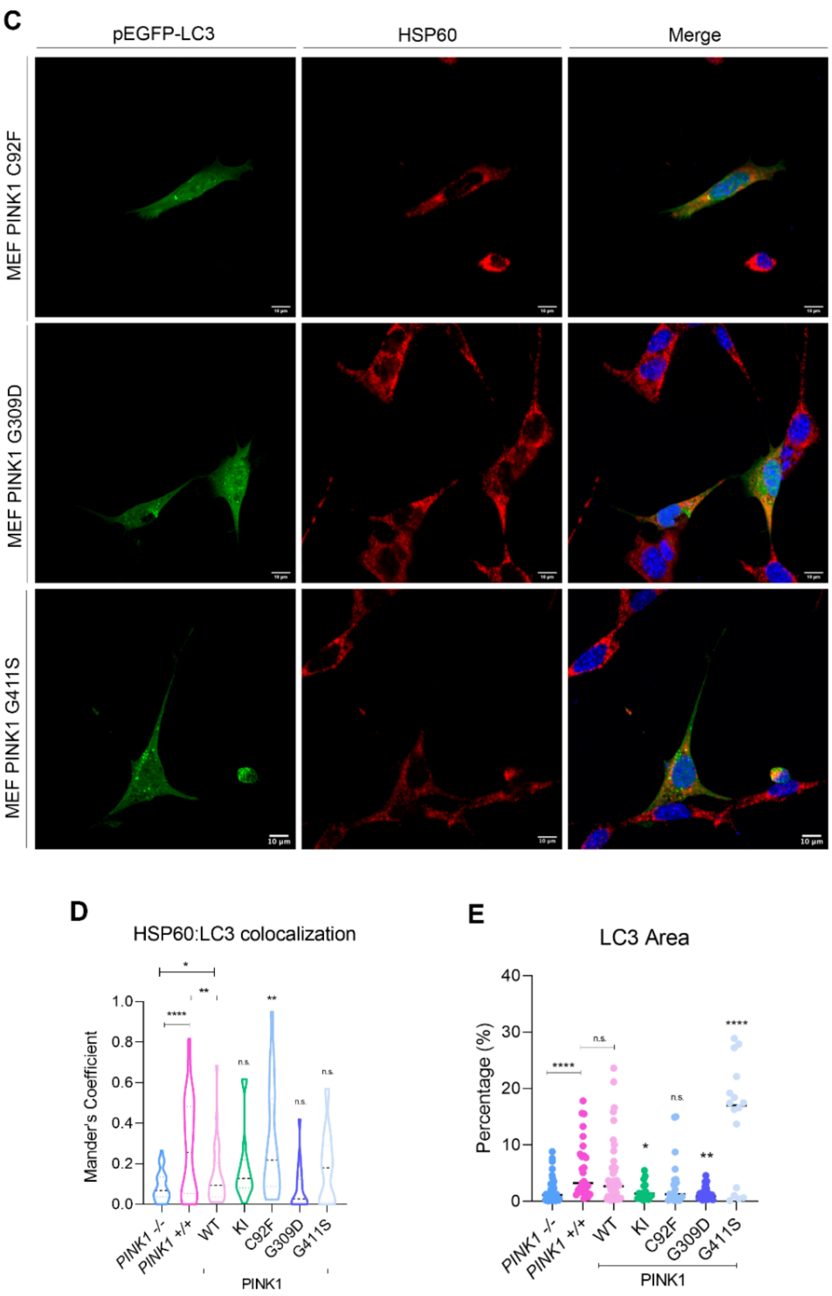


***Supplemental Figure S2** – Analysis of LC3 processing.*

*A) MEF transduced cell lines were treated with DMSO or 25µM CCCP and analyzed by SDS-PAGE on 4-15% Tris-acetate gel and further immunoblotted for PINK1, LC3 and GAPDH (as loading control). B-C) MEF transduced cell lines transfected with pEGFP-LC3 were depolarized with 25µM of CCCP (Scale bar=10µm). D) Colocalization of LC3B puncta with mitochondria (HSP60) was accessed in ImageJ plug-in JACoP which calculates Mander's Coefficient. E) LC3B puncta were count using ImageJ threshold tool analyzed, normalized to total cell area. Statistical significance was calculated using one way ANOVA followed by Dunnets test (* $P < 0.05$; ** $P < 0.01$; *** $P < 0.001$; **** $P < 0.0001$; n.s. – non significant).*

### Figure S3

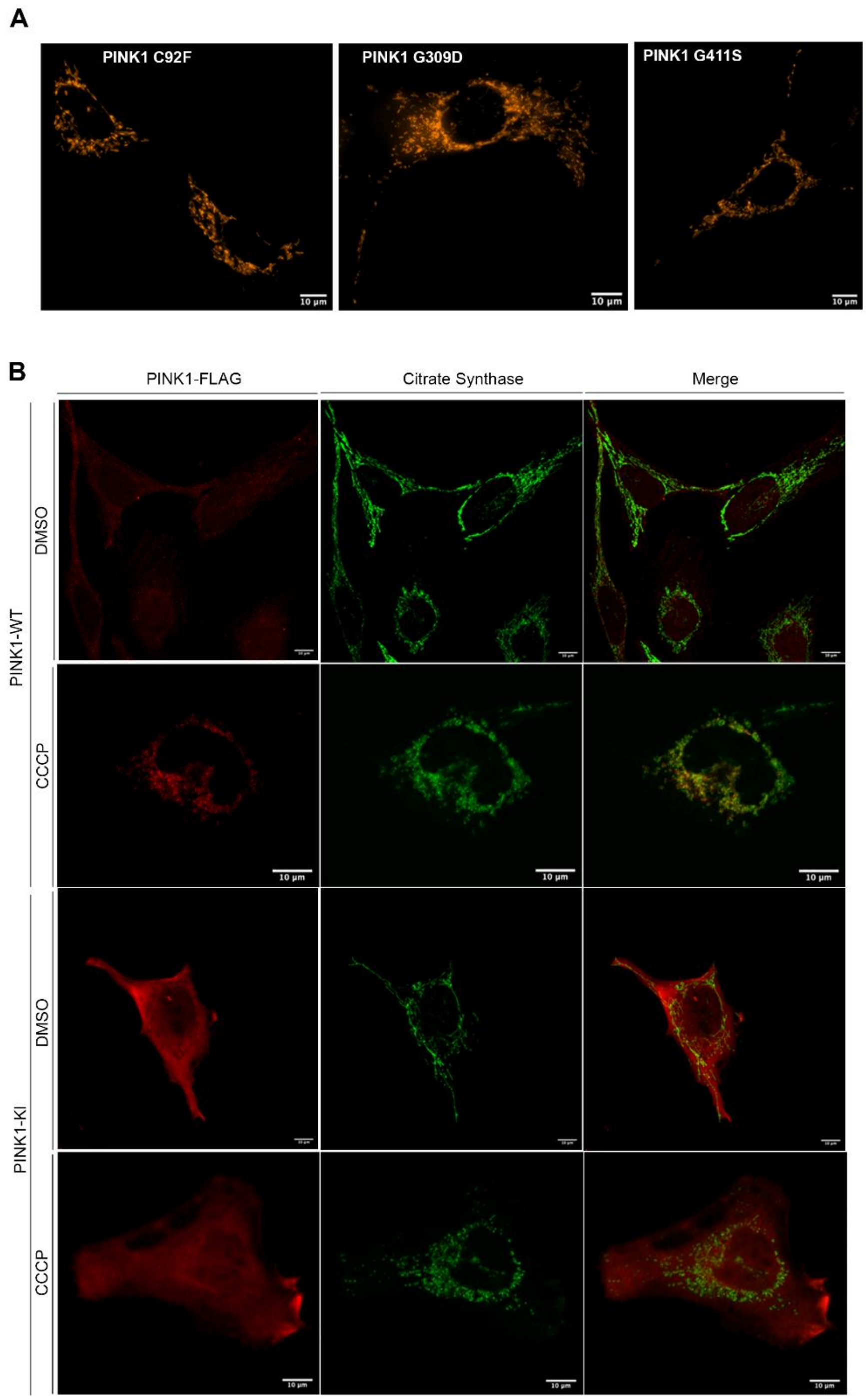
A
PINK1 C92F
PINK1 G309D
PINK1 G411S
10 µm
B
PINK1-FLAG
Citrate Synthase
Merge
PINK1-WT
DMSO
CCCP
PINK1-KI
DMSO
CCCP
10 µm

C

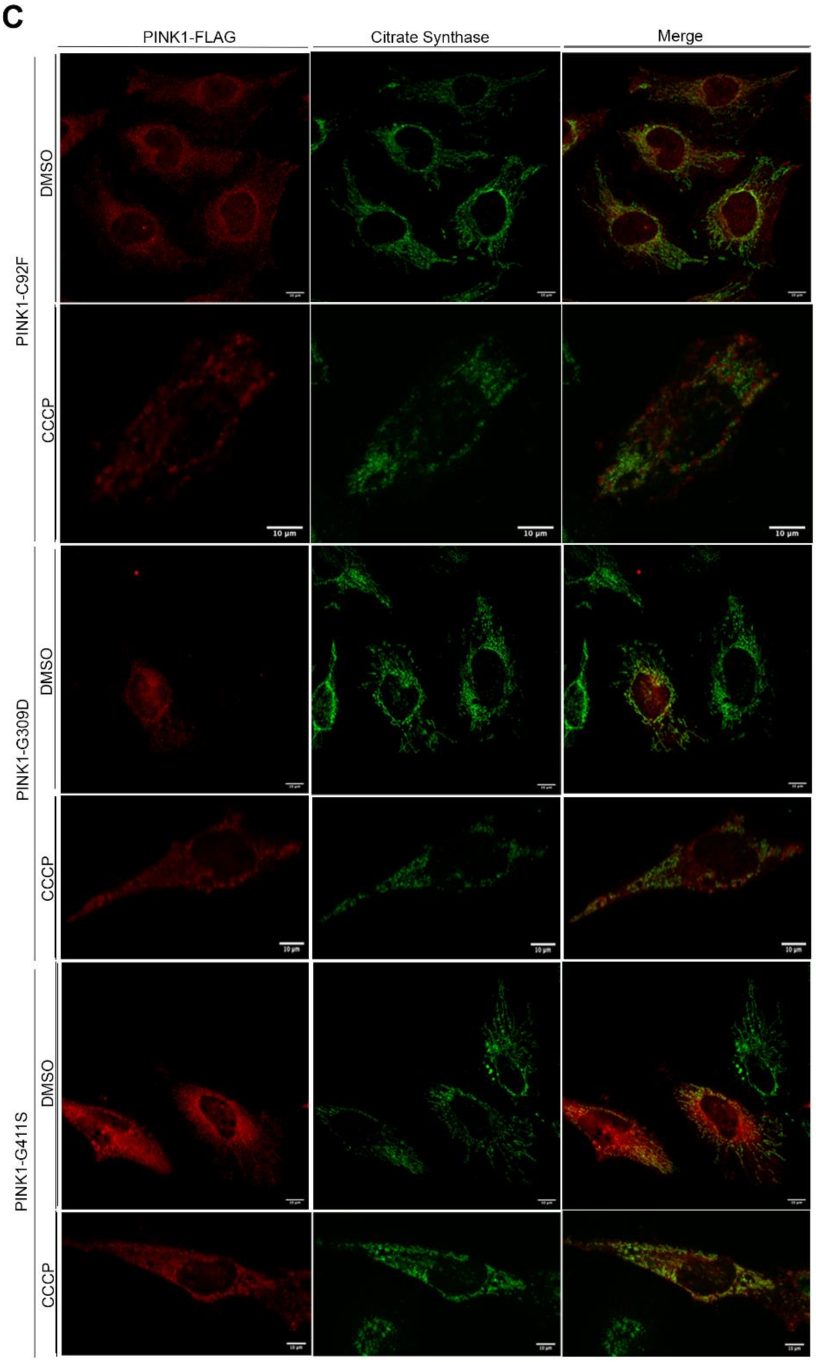


***Supplemental Figure S3*** *– Assessment of mitochondrial morphology in the presence of PINK1 mutants.*

*A) Mitochondrial morphology assessed in MEF cell line endogenously expressing PINK1 clinical mutations expressing a mitochondrial targeted mito-dsRed plasmid (Scale bar=10µm). B-C) HeLa PINK1$^{-/-}$ cells were transfected with FL-PINK1-WT, -KI and FLAG tagged clinical mutants and treated with 10µM CCCP. Citrate synthase staining (green) was used as a mitochondrial marker.*

**Figure S4**

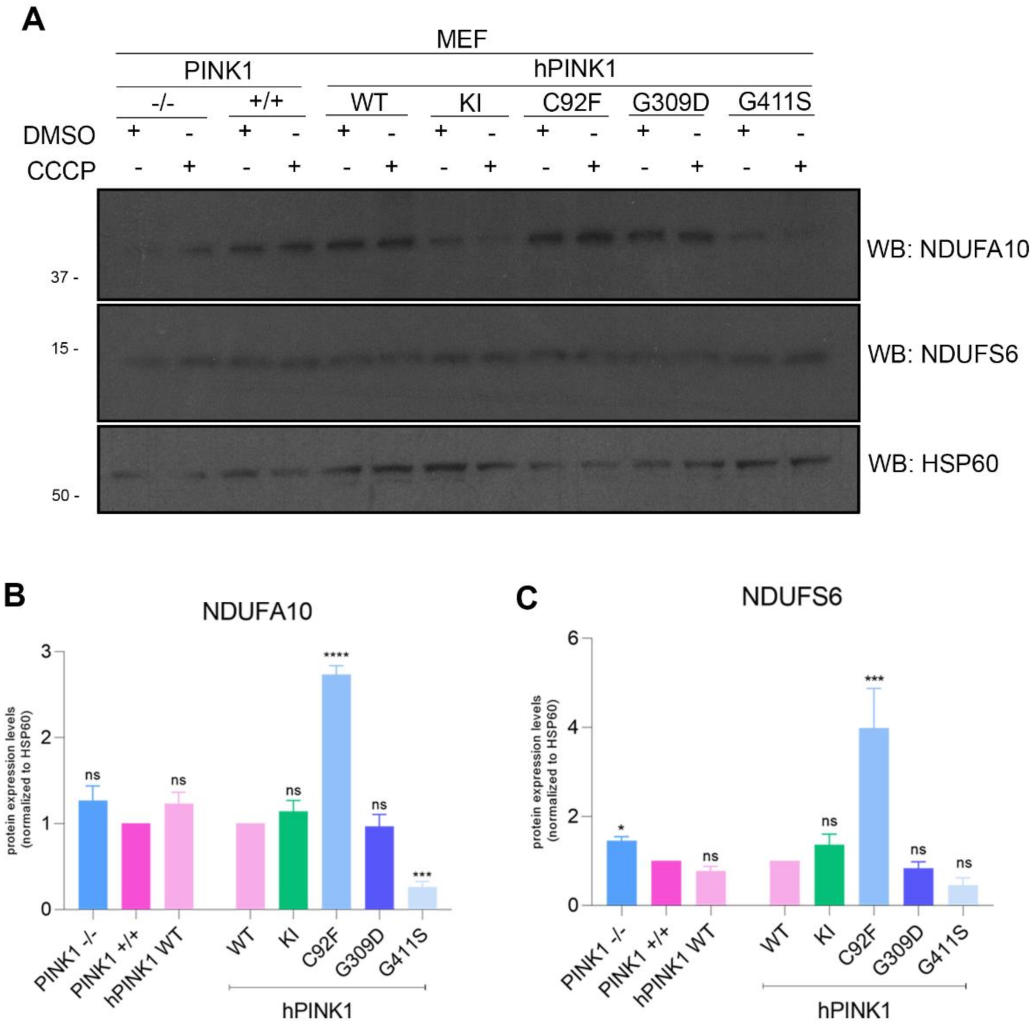


***Supplemental Figure S4*** *– Protein levels of Complex I subunits.*

*A) Western blot analysis and corresponding semi-quantification was performed to determine the expression levels of Complex I NDUFA10 (B) and NDUFS6 (C),. Statistical significance was calculated using one way ANOVA followed by Dunnets test (* P < 0.05; ** P < 0.01; *** P < 0.001; **** P < 0.0001; n.s. – non significant).*

**Figure S5**

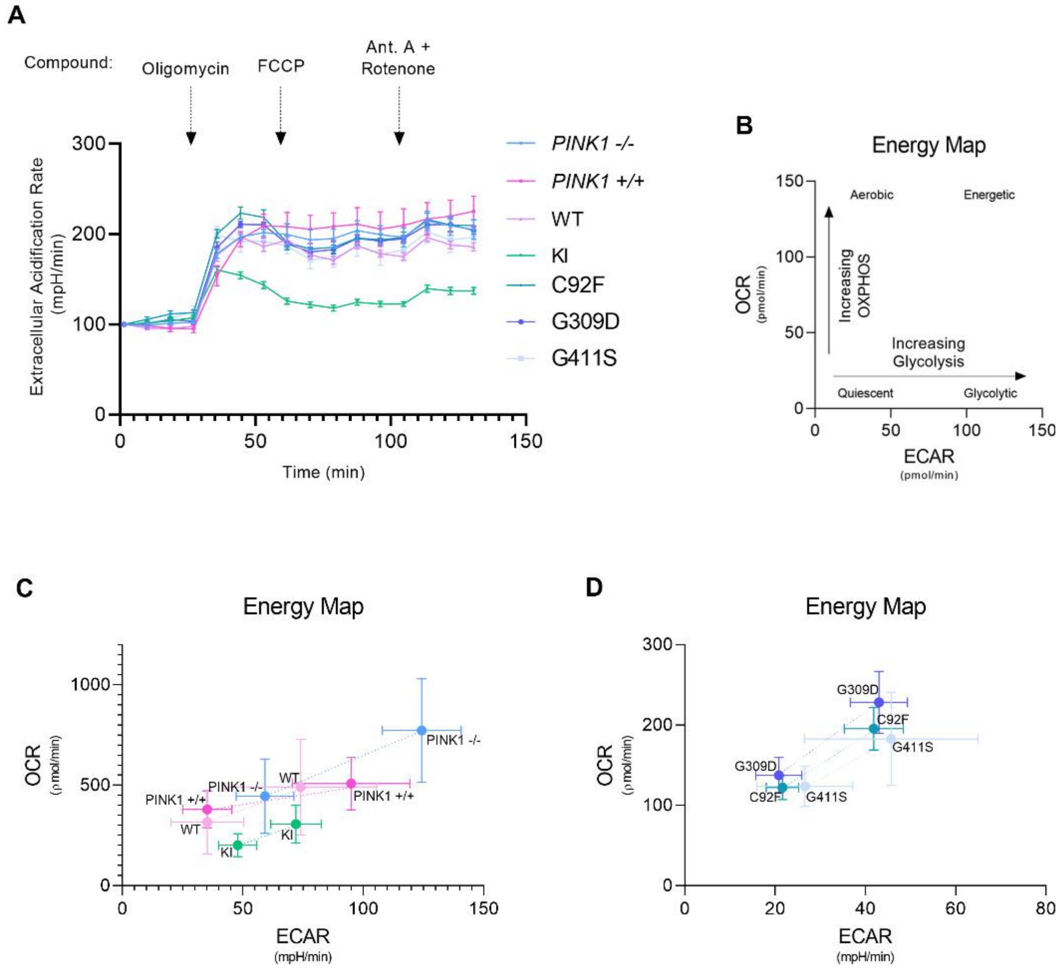


***Supplemental Figure S5** – Bioenergetic profiles of PINK1 mutants.*

*A) Extracellular acidification rate determined in MEF transduced cell lines injected with different compounds. B) Schematic representation of an Energy Map, where OCR is plotted with ECAR rising different energy cellular types. C) MEF PINK1-/-, MEF PINK1 +/+ and MEF hPINK1 WT and -KI cells plotted in an Energy Map. PINK1-/- have the better response when exposed in a stress condition. D) MEF hPINK1 transduced with -C92F, -G309D and -G411S plotted in an Energy Map, showing no major differences when in stress conditions.*

**Figure S6**

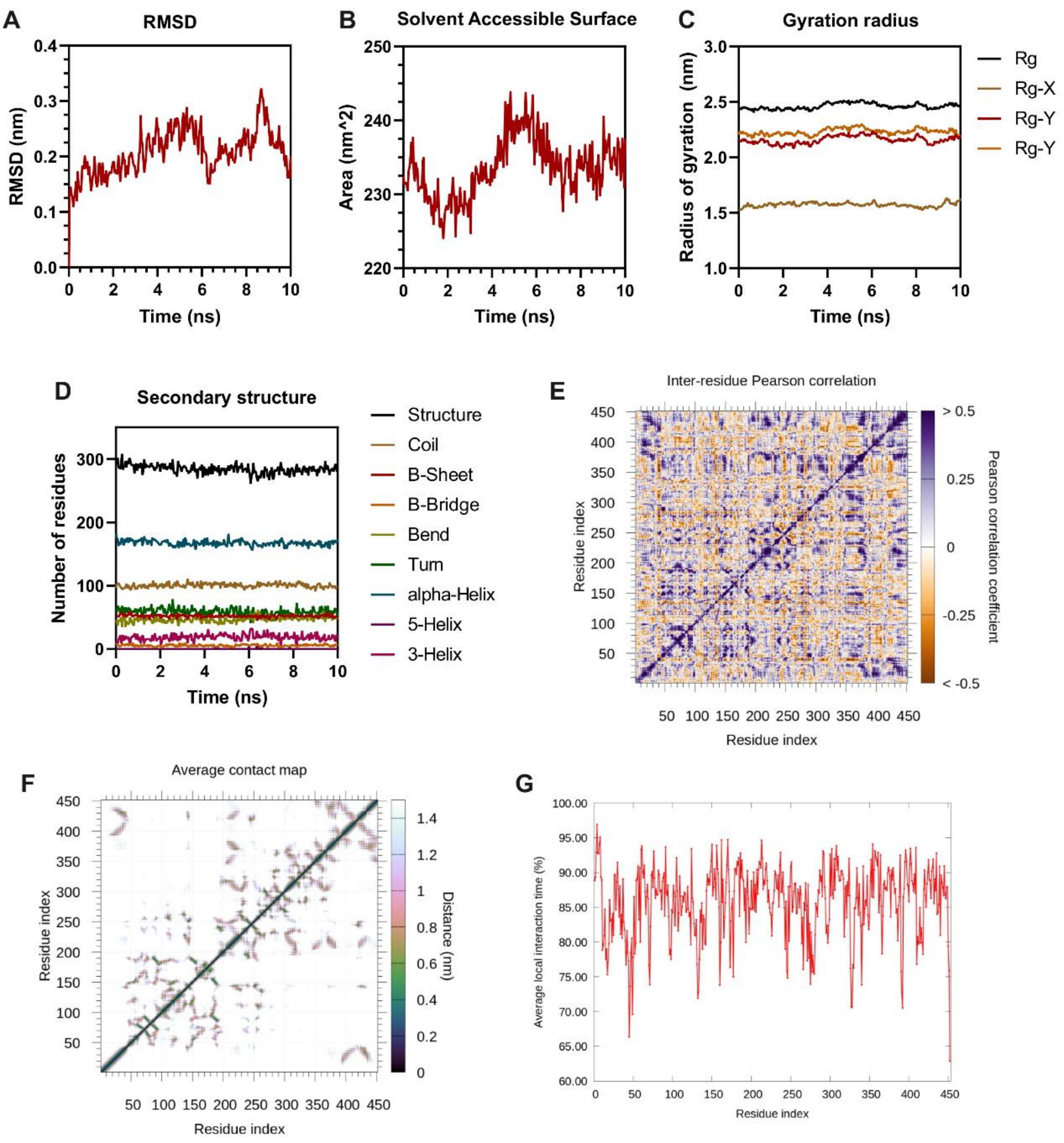


***Supplemental Figure S6** - Detailed analysis of the molecular dynamics simulations by GROMACS of the PINK1-WT protein structure.*

*A) Root mean square deviation, B) solvent accessible surface, C) gyration radius, D) evolution of secondary structure elements, E) inter-residue Pearson correlation, F) average contact map and G) average local interaction time along the protein structure.*

### Figure S7

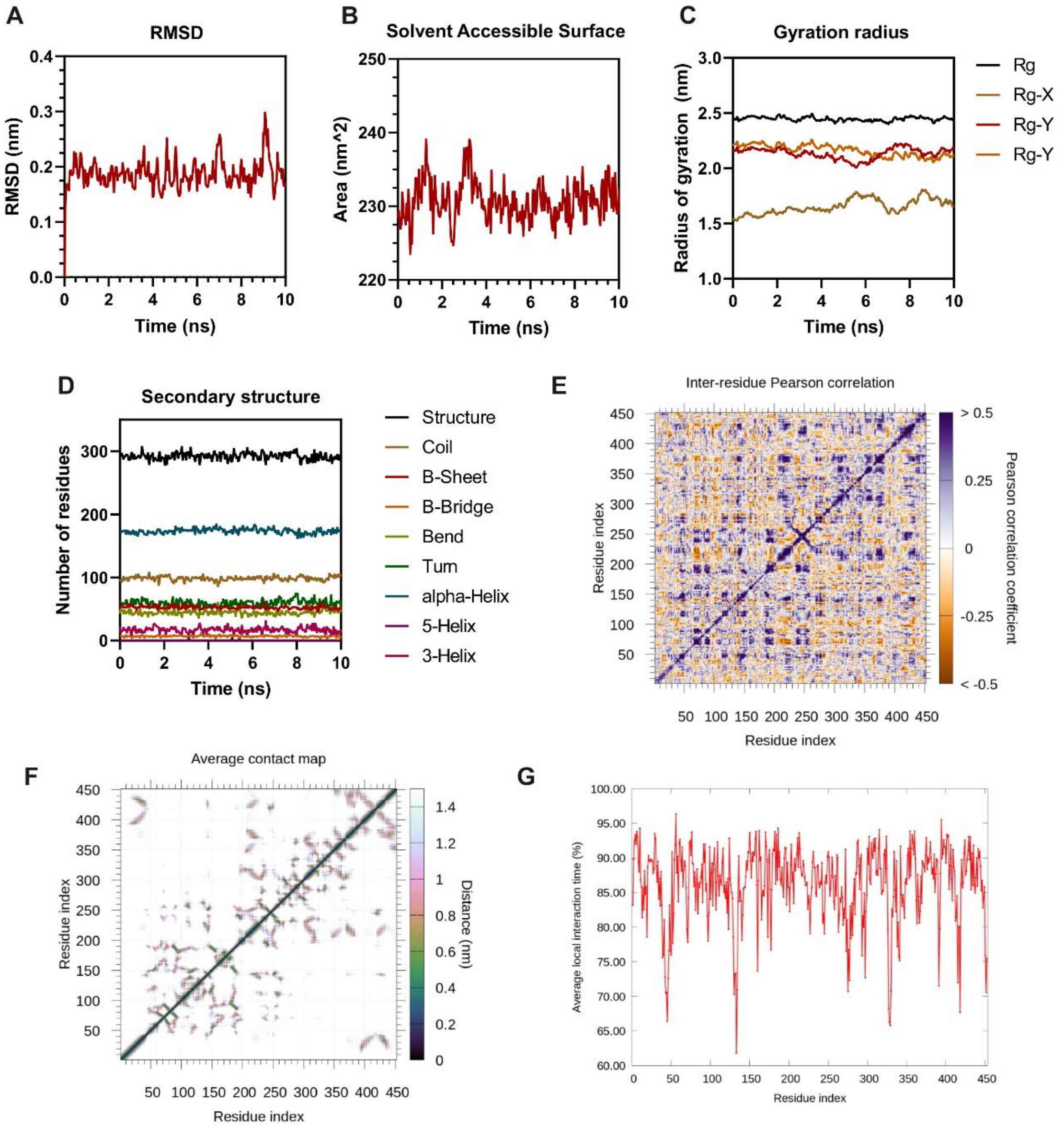


***Supplemental Figure S7*** *- Detailed analysis of the molecular dynamics simulations by GROMACS of the di-phosphorylated PINK1 protein structure at Serine228 and Serine402.*

*A) Root mean square deviation, B) solvent accessible surface, C) gyration radius, D) evolution of secondary structure elements, E) inter-residue Pearson correlation, F) average contact map and G) average local interaction time along the protein structure.*

**Figure S8**

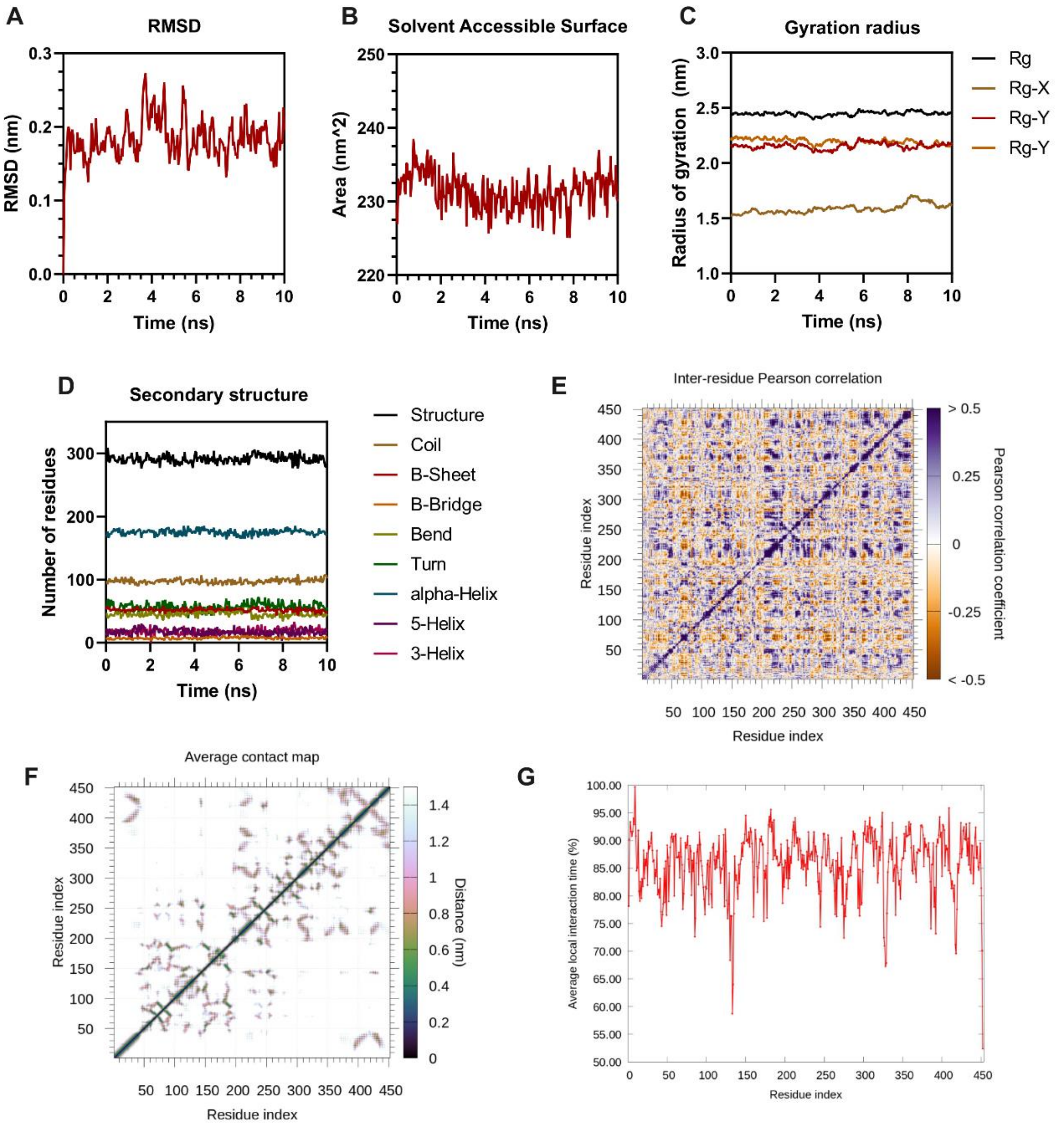


***Supplemental Figure S8*** *- Detailed analysis of the molecular dynamics simulations by GROMACS of the PINK1-G411S mutant structure.*

*A) Root mean square deviation, B) solvent accessible surface, C) gyration radius, D) evolution of secondary structure elements, E) inter-residue Pearson correlation, F) average contact map and G) average local interaction time along the protein structure.*

### Figure S9

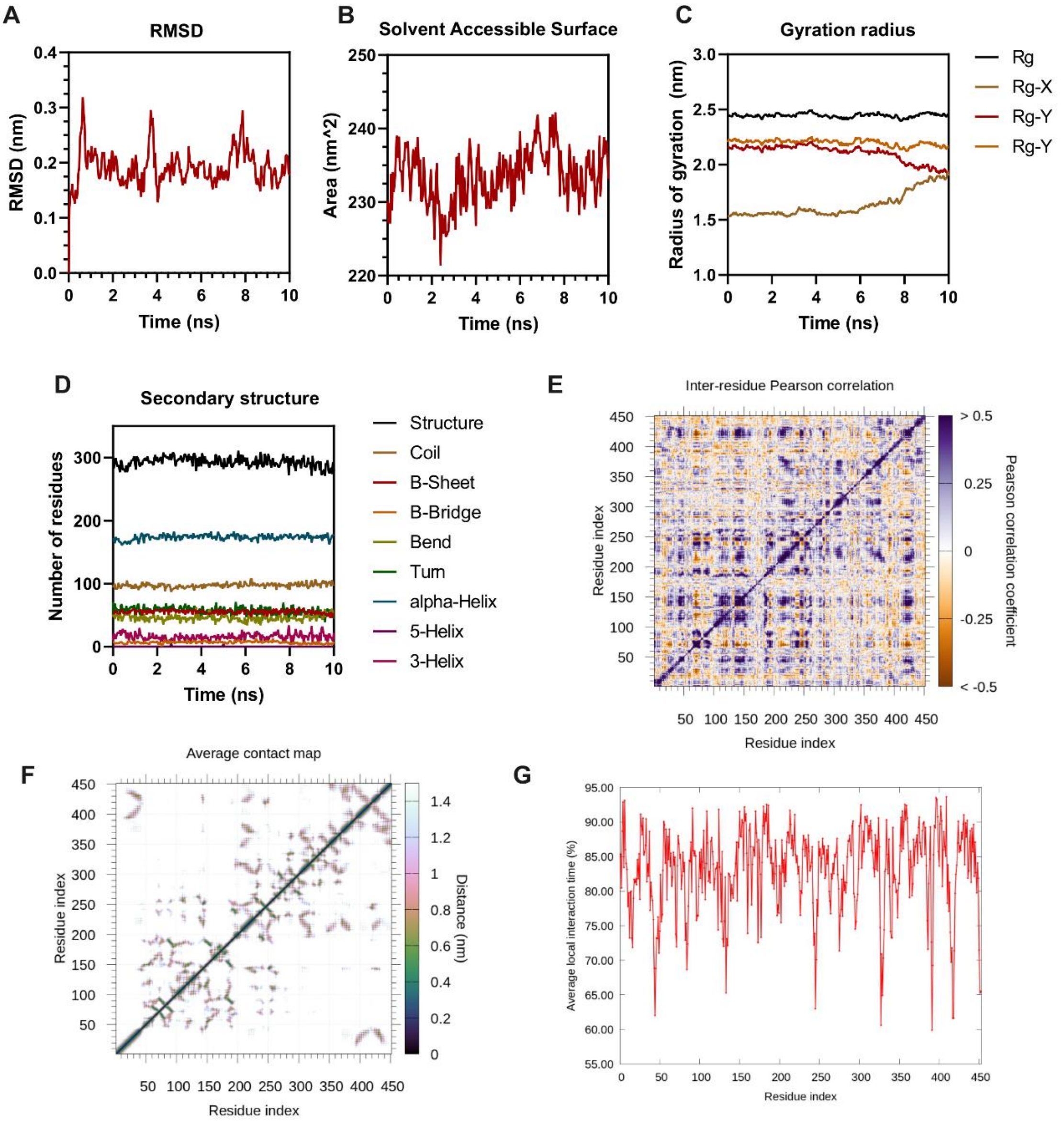


***Supplemental Figure S9*** *- Detailed analysis of the molecular dynamics simulations by GROMACS of the phosphorylated PINK1-G411S mutant protein structure at Ser411.*
*A) Root mean square deviation, B) solvent accessible surface, C) gyration radius, D) evolution of secondary structure elements, E) inter-residue Pearson correlation, F) average contact map and G) average local interaction time along the protein structure.*

**Figure S10**

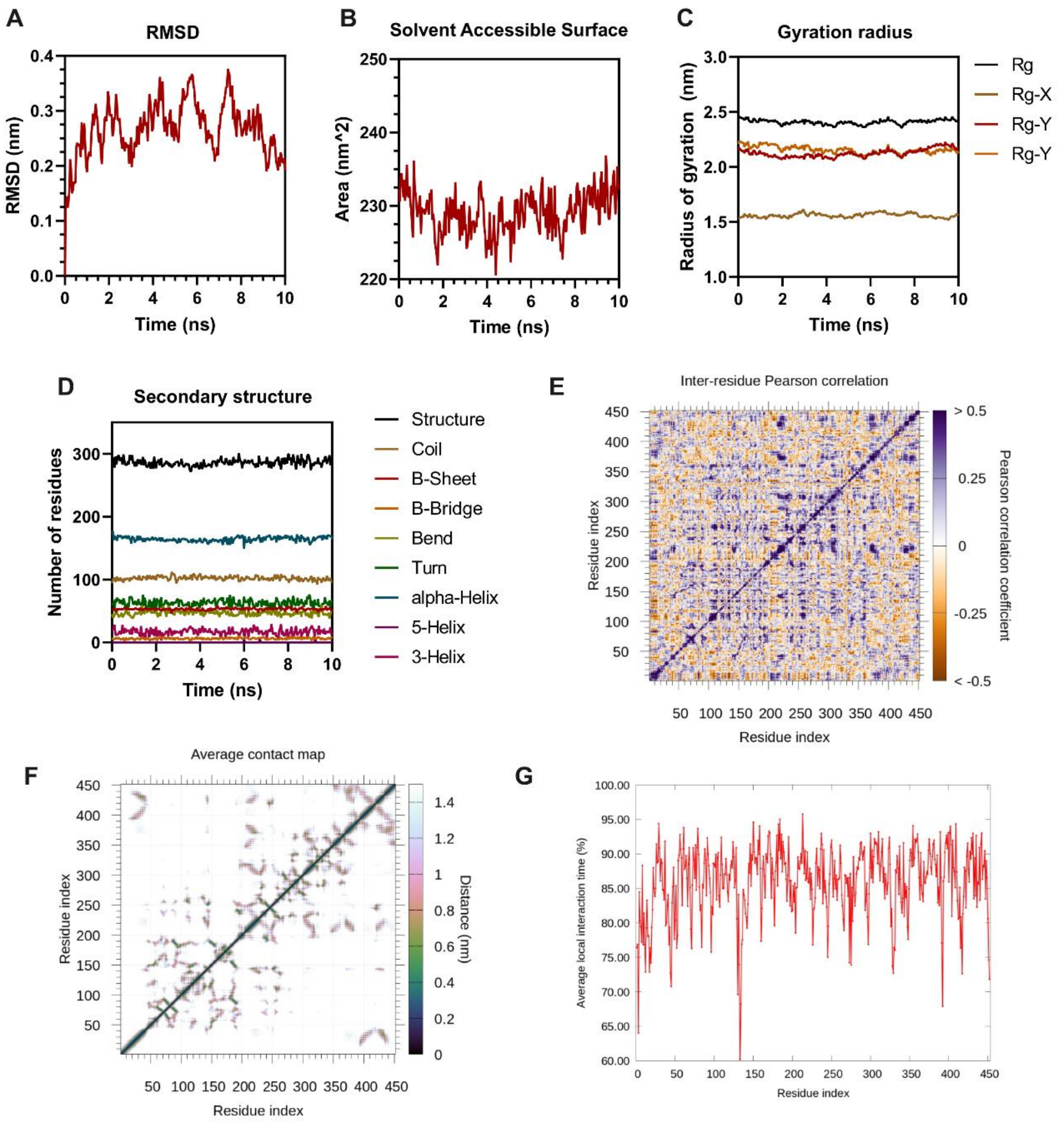


***Supplemental Figure S10*** *- Detailed analysis of the molecular dynamics simulations by GROMACS of the tri-phosphorylated PINK1-G411S mutant protein structure at Ser411, Serine228 and Serine402.*

*A) Root mean square deviation, B) solvent accessible surface, C) gyration radius, D) evolution of secondary structure elements, E) inter-residue Pearson correlation, F) average contact map and G) average local interaction time along the protein structure.*

***Table S1** – Selection of primers used for Quick Change Mutagenesis Protocol and Sequencing*

| | | | |
|---|---|---|---|
| **SEQUENCING PRIMERS** | hPINK1-middle | Forward | ACCTCTTCCGTGCCG |
| | | Reverse | CGGCACGGAAGAGGT |
| | pcDNA 3.1 – T7 | Forward | TAATACGACTCACTATAGGG |
| | pcDNA 3.1 – BGH | Reverse | TAGAAGGCACAGTCGAGG |
| **MUTAGENESIS PRIMERS** | hPINK1 C92F | Forward | CGGGCCTGGGGCTTCGCGGGC |
| | | Reverse | GCCCGCGAAGCCCCAGGCCCG |
| | hPINK1 A168P | Forward | GGTAAGGGCTGCAGTCCTGCTGTGTATGAAG |
| | | Reverse | CTTCATACACAGCAGGACTGCAGCCCTTACC |
| | hPINK1 H271Q | Forward | CCAAGCAACTAGCCCCTCAGCCCAACATCATC |
| | | Reverse | GCCGTGGACACCCCTGGGGCCATCA |
| | hPINK1 G309D | Forward | CTGAAGGCCTGGACCATGGCCGGAC |
| | | Reverse | GTCCGGCCATGGTCCAGGCCTTCAG |
| | hPINK1 L347P | Forward | CTGCTGCAGCTGCCGGAAGGCGTGGAC |
| | | Reverse | GTCCACGCCTTCCGGCAGCTGCAGCAG |
| | hPINK1 G411S | Forward | TCGGGGCGGAAACAGCTGTCTGATGGC |
| | | Reverse | GCCATCAGACAGCTGTTTCCGCCCCGA |
| | hPINK1 E417G | Forward | TGATGGCCCCAGGGGTGTCCACGGC |
| | | Reverse | GCCGTGGACACCCCTGGGGCCATCA |
| | hPINK1 L489P | Forward | TTGGTGAGGGCACCGCTCCAGCGAGAG |
| | | Reverse | CTCTCGCTGGAGCGGTGCCCTCACCAA |

***Table S2** – List of antibodies used for Western blotting and Immunofluorescence and their corresponding working dilution*

| Antibody | Company and Reference | Dilutions |
|---|---|---|
| **Monoclonal ANTI-FLAG® M2 Antibody produced in mouse** | Sigma-Aldrich #F3165-5MG | 1/500 |
| **Anti-Glutathione-S-Transferase (GST) Antibody produced in rabbit** | Sigma-Aldrich #G7781-100UL | 1/2000 |
| **Purified Mouse Anti-Hsp60 Antibody** | BD Biosciences #611563 | 1/1000 |
| **Anti-Turbo GFP Antibody** | Evrogen #ab514 | 1/1000 |
| **anti-LC3 Antibody produced in Rabbit** | MBL International #PM036 | 1/1000 |
| **GAPDH Monoclonal Antibody** | Alfagene - Thermo Fisher Scientific - Life technologies #AM4300 | 1/5000 |
| **PINK1 polyclonal Antibody** | Novus Biologicals #BC100-494 | 1/1000 |
| **Total OXPHOS Rodent WB Antibody Cocktail** | Abcam #ab110413 | 1/5000 |
| **Citrate Synthase Antibody** | Proteintech #10837-1-AP | 1/250 |
| **Goat Anti-Mouse HRP Conjugated Secondary Antibody** | Bio-Rad #1706516 | 1/10000 |
| **Goat Anti-Rabbit HRP Conjugated Secondary Antibody** | Bio-Rad #1706515 | 1/10000 |
| **Alexa 488 Donkey anti-RABBIT IgG (H+L) Secondary Antibody** | Molecular Probes - Thermo Fisher #A21206 | 1/500 |
| **Alexa 568 goat anti-MOUSE IgG (H+L) Secondary Antibody** | Molecular Probes - Thermo Fisher #A11031 | 1/500 |
